# Evolution of wheat blast resistance gene *Rmg8* accompanied by differentiation of variants recognizing the powdery mildew fungus

**DOI:** 10.1101/2023.09.26.559445

**Authors:** Soichiro Asuke, Kohei Morita, Motoki Shimizu, Fumitaka Abe, Ryohei Terauchi, Chika Nago, Yoshino Takahashi, Mai Shibata, Motohiro Yoshioka, Mizuki Iwakawa, Mitsuko Kishi-Kaboshi, Zhuo Su, Shuhei Nasuda, Hirokazu Handa, Masaya Fujita, Makoto Tougou, Koichi Hatta, Naoki Mori, Yoshihiro Matsuoka, Kenji Kato, Yukio Tosa

## Abstract

Wheat blast, a devastating disease having spread recently from South America to Asia and Africa, is caused by *Pyricularia oryzae* pathotype *Triticum* which emerged in 1985. *Rmg8* and *Rmg7*, genes for resistance to wheat blast found in common wheat and tetraploid wheat, respectively, recognize the same avirulence gene, *AVR-Rmg8*. Here, we show an evolutionary process in which resistance gene(s), which had obtained an ability to recognize *AVR-Rmg8* before the differentiation of *Triticum* and *Aegilops*, has expanded its target pathogens. Molecular cloning revealed that *Rmg7* was one of alleles of *Pm4* (*Pm4a*), a gene for resistance to wheat powdery mildew on 2AL, whereas *Rmg8* was its homoeolog on 2BL ineffective against wheat powdery mildew. *Rmg8* variants with the ability to recognize *AVR-Rmg8* were distributed not only in *Triticum* spp. but also in *Aegilops speltoides*, *Ae. umbellulata,* and *Ae. comosa*. This result suggests that the origin of resistance gene(s) recognizing *AVR-Rmg8* dates back to the time before differentiation of A, B, S, U, and M genomes, that is, ∼5 million years before the emergence of its current target, the wheat blast fungus. Phylogenetic analyses suggested that, in the evolutionary process thereafter, some of their variants gained the ability to recognize the wheat powdery mildew fungus and evolved into genes for resistance to wheat powdery mildew.

## Introduction

Wheat cultivation is now threatened by an expanding pandemic disease – wheat blast^1^. Its causal agent is a subgroup of a filamentous fungus, *Pyricularia oryzae* (syn. *Magnaporthe oryzae*) pathotype *Triticum* (MoT)^2^, which is specifically pathogenic on the genus *Triticum*^3^. MoT first emerged in Brazil in 1985^4^ through a host jump of *P. oryzae* pathotype *Lolium* (MoL) or its relatives^5^, then spread to neighboring countries such as Bolivia, Paraguay, and Argentina, and became one of the most serious wheat diseases in South America. Recently, it spread to Asia and Africa, and caused severe outbreaks of wheat blast in Bangladesh (in 2016)^6–8^ and Zambia (in 2018)^9^. Molecular analyses of isolates collected in these countries suggested that the outbreaks in Bangladesh and Zambia were caused by a lineage which spread from South America to Asia and Africa through independent introductions^1^. To control this devastating disease, we need resistance genes effective against MoT. The only genetic resource currently used in farmer’s field against MoT is a 2NS chromosomal segment^10^ introduced from *Aegilops ventricosa*^11^. However, the resistance gene on this segment has not been identified. Furthermore, the 2NS resistance has already been overcome by new MoT strains in South America^10,12^.

Genes for resistance to MoT have been considered to be rarely found in the current wheat population because MoT is a new pathogen which emerged only ∼40 years ago; most of current wheat accessions have not been exposed to the attack or infection pressures by MoT. However, Tagle et al.^13^ identified a resistance gene in cultivated emmer wheat and designated it as *Rmg7*. Anh et al.^14^ identified another resistance gene in common wheat cultivar S-615 and designated it as *Rmg8*. *Rmg7* and *Rmg8* were located on distal ends of the long arms of chromosome 2A (2AL) and 2B (2BL), respectively^14^. In addition, these genes corresponded to the same avirulence gene, *AVR-Rmg8*^15^. These results suggested that they might be homoeologous genes derived from the same ancestral gene^15^. To be useful in farmer’s fields, wheat blast resistance genes must be effective even at high temperature because wheat blast is severe at high temperature with an optimum between 25 and 30℃^2^. *Rmg8* was effective at high temperature but *Rmg7* was not^15^, suggesting that *Rmg8* is the only wheat blast resistance gene identified so far as a major gene which may be useful in fields.

To find additional resistance genes, Wang et al.^16^ screened 520 local landraces of common wheat collected from various countries over the world, and found 18 accessions resistant to MoT. We initially expected that several resistance genes might be found in these resistant accessions. However, all of these accessions recognized *AVR-Rmg8*, which led us to infer that they are all *Rmg8* carriers although one of them had an additional gene tentatively designated as *RmgGR119*^16^. These resistant accessions had been collected in Europe and Middle East between 1924 and 1971 (before the emergence of MoT), and should not have had interactions with MoT. Why have these accessions in Europe and Middle East maintained *Rmg8*, a gene for resistance to MoT? In the present study we isolated *Rmg8* and *Rmg7*, and found that they are a homoeolog and an allele, respectively, of *Pm4*, a gene for resistance to wheat powdery mildew (*Blumeria graminis* f. sp. *tritici*, Bgt). In addition, we found their functional variants in *Aegilops speltoides*, *Ae. umbellulata*, and *Ae. comosa*, suggesting that the origin of resistance genes recognizing *AVR-Rmg8* dates back to the time before the differentiation of *Triticum* and *Aegilops*, that is, ∼5 million years before the emergence of its target pathogen, MoT. Based on these results, we present a model of evolutionary processes in which a resistance gene has gained new target pathogens through differentiation of variants.

## Results

### *Rmg8* is a homoeolog of *Pm4*, a gene for resistance to wheat powdery mildew

To isolate *Rmg8* from common wheat, the resistant cultivar S-615 carrying *Rmg8* was crossed with a susceptible cultivar, Shin-chunaga (Sch), resulting in 165 F_2:3_ lines. When inoculated with MoT isolate Br48, homozygous resistant, segregating, and homozygous susceptible lines segregated in a 1:2:1: ratio (45:87:33) as expected. Molecular markers (KM markers) for mapping were produced using high confidence genes on 2BL found in the whole genome sequence of cv. Chinese Spring in the database (International Wheat Genome Sequencing Consortium; IWGSC). Mapping with the KM markers delimited the candidate region to ∼12Mb between KM25 and the telomere (Fig. 1a). However, we could not narrow down the candidate region further because all markers produced on its 8.4Mb distal region co-segregated with the *Rmg8* phenotype (Fig. 1a).

**Fig. 1.**
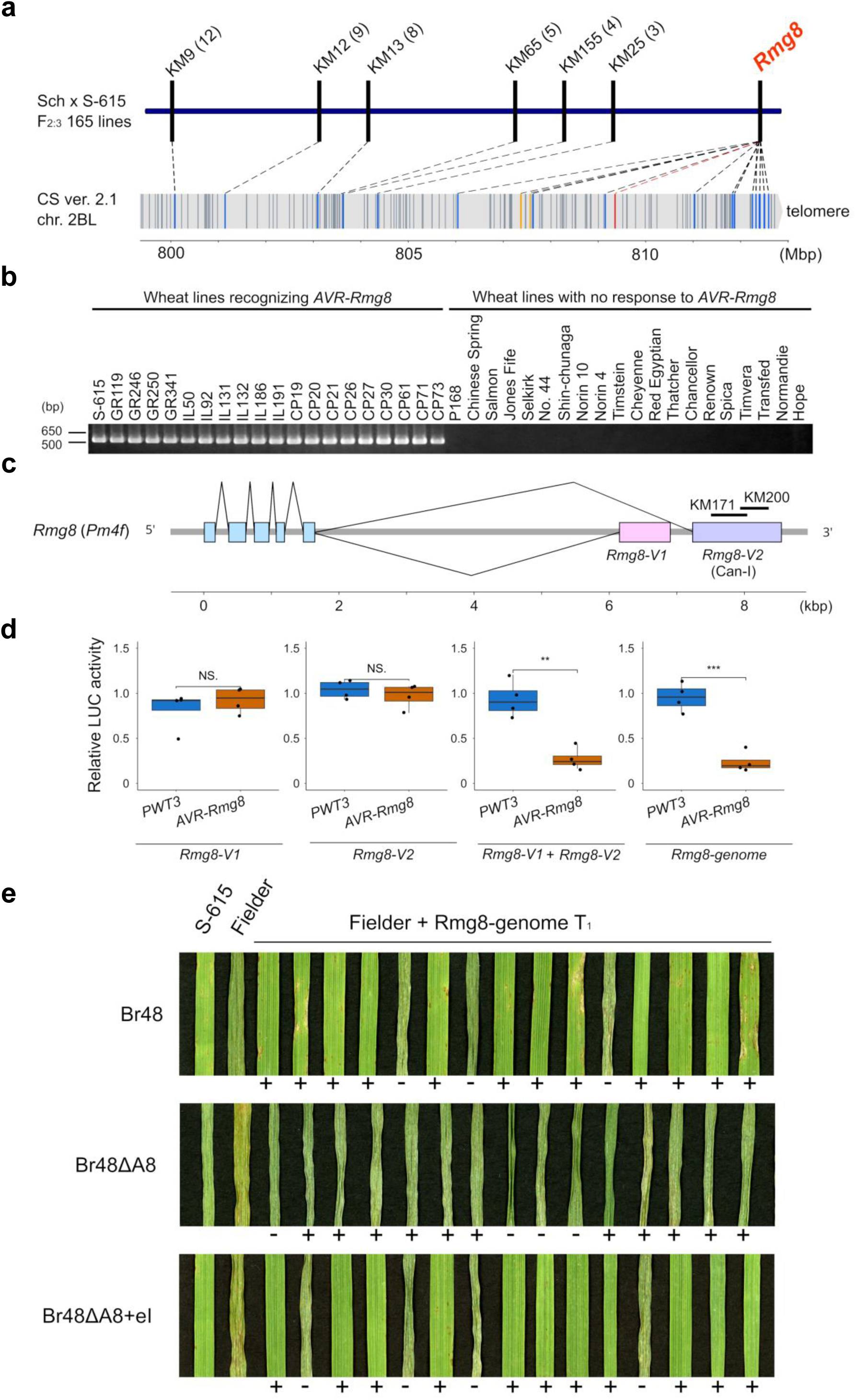
Cloning of *Rmg8* identified on chr. 2BL in common wheat. **a**, Genetic and physical maps around *Rmg8* on chr. 2BL. In the genetic map, numbers of recombinants between each molecular marker and *Rmg8* are shown in parentheses. In the physical map, positions of high confidence genes annotated in the Chinese Spring reference genome v2.1 are indicated by vertical lines. Genes used as molecular markers, a Can-I-like gene, and other genes picked up through the association analysis (see Extended Data Fig. 1) are highlighted with blue, red, and yellow, respectively. **b**, Association between responses to *AVR-Rmg8* and amplifications with the KM171 primers in common wheat. **c**, Structure of the gene producing the Can-I transcript. Can-I was one of the two splicing variants (*Rmg8-V1* and *Rmg8-V2*) of the gene. Bold lines represent positions of presence/absence PCR markers (KM171 and KM200) used for mapping and detection of *Rmg8*. **d**, Cell death assay with protoplasts. Protoplasts isolated from barley primary leaves were transfected with pAHC17-LUC containing a luciferase gene, pZH2Bik containing avirulence genes (*PWT3* or *AVR-Rmg8* lacking signal peptides) or no insert (empty vector), and pZH2Bik containing constructs of the *Rmg8* candidate gene (*Rmg8-V1*, *Rmg8-V2*, *Rmg8-genome*) or a mixture of pRmg8-V1 and pRmg8-V2 in a 1:1 molar ratio. Luciferase activity was determined 18-hours after transfection and represented as relative activities compared with those in samples with the empty vector. Double and triple asterisks indicate significant differences at the 1 and 0.1% levels, respectively, in the Tukey post hoc test. NS, not significant. The experiments were repeated four times. **e**. Validation of *Rmg8* through transformation. S-615 (*Rmg8*), Fielder (*rmg8*), and T_1_ individuals derived from transformation of Fielder with the genomic fragment (*Rmg8-genome*) were inoculated with Br48 (wild MoT isolate), Br48ΔA8 (disruptant of *AVR-Rmg8* derived from Br48), and Br48ΔA8+eI (transformant of Br48ΔA8 carrying re-introduced *AVR-Rmg8* derived from Br48), and incubated for five days. Presence (+) /absence (-) of the transgene confirmed by PCR with the KM171 and HPT markers are shown below the panels.

We then adopted the RaIDeN method developed by Shimizu et al.^17^ with some modifications. Briefly, RNA-seq reads obtained from primary leaves of Sch and nine F_2:3_ lines with homozygous susceptible genotypes were aligned to a reference sequence of a gene set which was constructed by de novo assembly of RNA-seq reads obtained from S-615 leaves (Extended Data Fig. 1a). We selected genes (i) which showed polymorphisms (presence/absence or single-nucleotide polymorphisms) between S-615 and Sch, (ii) whose Sch allele was shared by all of the nine susceptible F_2:3_ lines, and (iii) which encoded NBS (nucleotide-binding site), NLR (nucleotide-binding site – leucine-rich repeat), or RLK (receptor-like kinase). Consequently we found 10 genes that fulfilled the three requirements (Extended Data Fig. 1b). *In silico* analyses with whole genome sequences in the databases suggested that six out of the 10 genes were located on the 2B chromosome of *T. turgidum* subsp. *durum* cv. Svevo. PCR markers designed on these 6 genes co-segregated with *Rmg8* in the F_2:3_ lines derived from Sch x S-615, indicating that they are actually located on 2B of common wheat. Finally, these candidate genes were subjected to an association analysis using 20 common wheat lines recognizing *AVR-Rmg8* (S-615, the 18 local landraces mentioned above, and GR341, an additional local landrace which proved to recognize *AVR-Rmg8*) and 20 common wheat cultivars that did not recognize *AVR-Rmg8*. One candidate, the Can-I gene amplified with PCR marker KM171, showed a perfect association with the *Rmg8* phenotype (Fig. 1b) whereas the others did not (Extended Data Fig. 1b). From these results, we assumed that the Can-I gene might be *Rmg8*. Intriguingly, the Can-I transcript sequence was almost identical to that of *Pm4b*_*V2*, one of the two splicing variants of *Pm4b* controlling the resistance to wheat powdery mildew^18^. Further analyses revealed that transcripts from S-615 also contained another splicing variant which was almost identical to *Pm4b_V1*, the other splicing variant of *Pm4b*. These alternative splicing variants derived from S-615 were designated as *Rmg8-V2* and *Rmg8-V1*, respectively. A comparison of these transcripts with the genome sequence of S-615 revealed that the exon/intron structure was the same as *Pm4b* (Fig. 1c).

To check whether *Rmg8-V1* and *Rmg8-V2* recognize *AVR-Rmg8* and induce hypersensitive reaction, a protoplast cell death assay^19^ was performed. cDNA fragments of *Rmg8-V1*, *Rmg8-V2*, and a genomic fragment of the entire gene was inserted into pZH2Bik^20^ so as to be driven by the rice ubiquitin promoter, and established as pRmg8-V1, pRmg8-V2, and pRmg8-genome, respectively. Barley protoplasts were co-transfected with these constructs, a plasmid carrying *AVR-Rmg8* (pAVR-Rmg8), and a plasmid carrying a luciferase reporter gene. As a negative control, pPWT3 with the *PWT3* avirulence gene^5^ corresponding to *Rwt3* was employed instead of pAVR-Rmg8. Fluorescence was not reduced in pAVR-Rmg8/pRmg8-V1 or pAVR-Rmg8/pRmg8-V2 combinations (Fig. 1d). When protoplasts were co-transfected with pAVR-Rmg8 and a mixture of pRmg8-V1 and pRmg8-V2, however, fluorescence was significantly reduced. This reduction was also observed in the pAVR-Rmg8/pRmg8-genome combination, but was cancelled when pAVR-Rmg8 was replaced with pPWT3. These results suggest that the Can-I gene is *Rmg8* and that both of its splicing variants are required for the recognition of *AVR-Rmg8*. This is in accordance with the report^18^ that both of *Pm4b_V1* and *Pm4b_V2* are required for the resistance to powdery mildew conferred by *Pm4b*.

To confirm that the Can-I gene is *Rmg8*, pRmg8-genome was introduced into *T. aestivum* cv. Fielder (susceptible to Br48) through Agrobacterium-mediated transformation. In the T_1_ generation, resistant and susceptible individuals against Br48 segregated in a 3:1 ratio (Fig. 1e). Furthermore, these reactions to Br48 were perfectly concordant with the presence/absence of the transgene. By contrast, the T_1_ individuals were all susceptible to Br48ΔA8 (*AVR-Rmg8* disruptant derived from Br48) irrespective of the presence/absence of the transgene. Against Br48ΔA8+eI (transformant of Br48ΔA8 carrying re-introduced *AVR-Rmg8* derived from Br48), resistant and susceptible T_1_ individuals again segregated in a 3:1 ratio in concordance with the presence/absence of the transgene. Transformants carrying pRmg8-V1 alone and those carrying pRmg8-V2 alone were all susceptible to Br48, Br48ΔA8, and Br48ΔA8+eI (Extended Data Fig. 2), supporting the observation in the protoplast assay. From these results, we concluded that we successfully isolated *Rmg8*. Sánchez-Martin et al.^18^ found six “alleles” of *Pm4*, i.e., *Pm4a*, *Pm4b*, *Pm4d*, *Pm4f*, *Pm4g*, *Pm4h* in breeding lines or global collections of common wheat through PCR amplification and Sanger sequencing. The genetically identified *Pm4* alleles, i.e., *Pm4a*, *Pm4b*, and *Pm4d*, were located on 2A^21,22^ while chromosomal locations of *Pm4f*, *Pm4g*, *Pm4h* have not been determined. *Rmg8* was identical to *Pm4f* in the nucleotide sequence, but was located on 2B (Fig. 2a). From these results, we concluded that *Rmg8* is a homoeolog of *Pm4*.

**Fig. 2.**
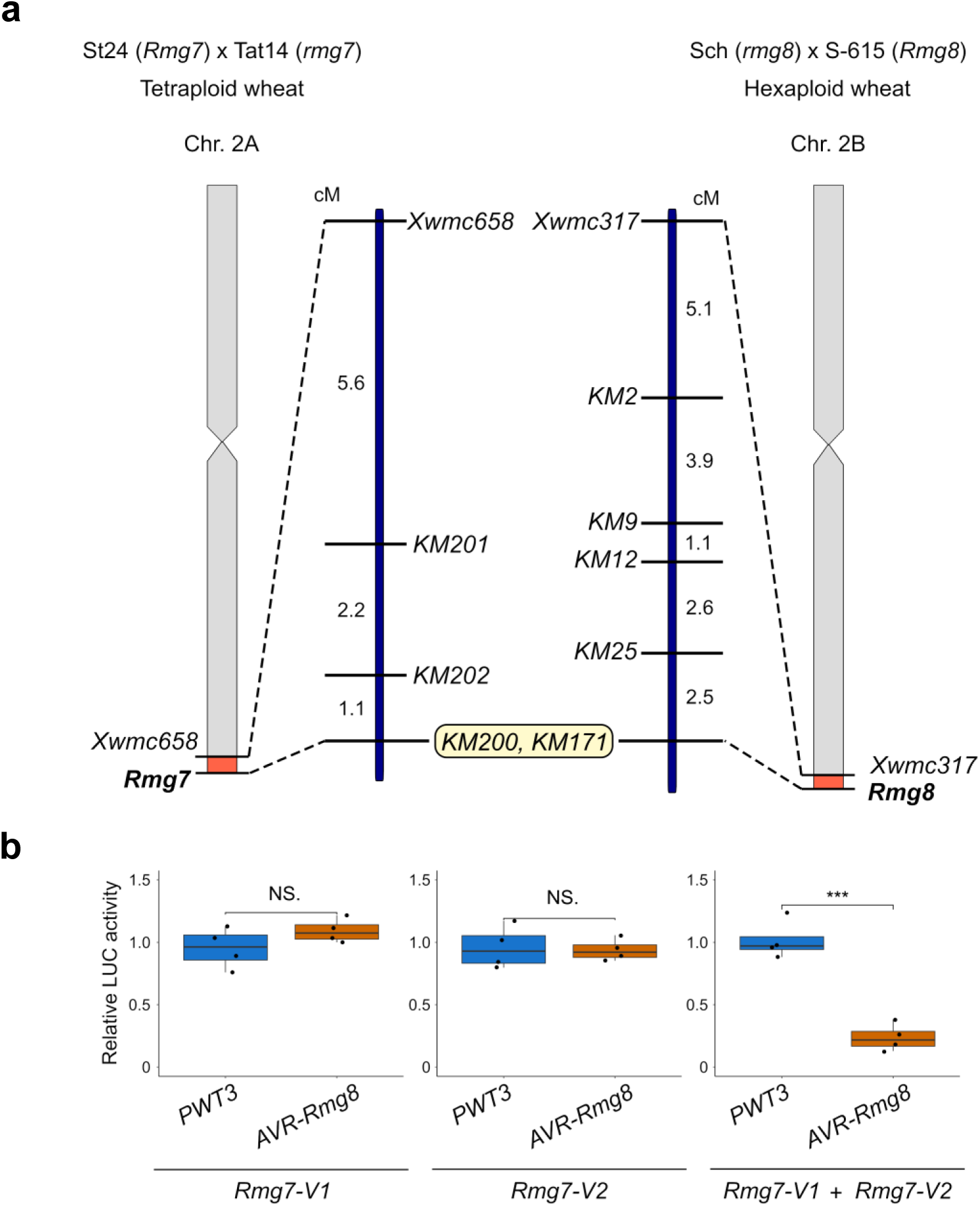
Cloning of *Rmg7* identified on chr. 2AL in tetraploid wheat. **a**, Genetic map around *Rmg7* constructed using 93 F_2:3_ lines derived from *T. dicoccum* accession St24 (*Rmg7*) x *T. paleocolchicum* accession Tat14 (*rmg7*). For a comparison a genetic map around *Rmg8* on 2BL was shown on the right which was constructed using 91 F_2:3_ lines derived from *T. aestivum* cv. Shin-Chunaga (Sch, *rmg8*) x *T. aestivum* cv. S-615 (*Rmg8*). KM200 and KM171, PCR markers designed on *Rmg8*, perfectly co-segregated with phenotypes conferred by *Rmg7*. **b**, Cell death assay with protoplasts. Protoplasts isolated from barley primary leaves were transfected with pAHC17-LUC containing a luciferase gene, pZH2Bik containing avirulence genes (*PWT3* or *AVR-Rmg8* lacking signal peptides) or no insert (empty vector), and pZH2Bik containing constructs of the *Rmg7* candidate gene (Rmg7-V1, Rmg7-V2) or a mixture of pRmg7-V1 and pRmg7-V2 in a 1:1 molar ratio. Luciferase activity was determined 18-hours after transfection and represented as relative activities compared with those in samples with the empty vector. Triple asterisks indicate significant differences at the 0.1 % level in the Tukey post hoc test. NS, not significant. The experiments were repeated four times.

### *Rmg7* is an allele of *Pm4*, a gene for resistance to wheat powdery mildew

*Rmg7* was identified in three accessions of tetraploid wheat, *T. dicoccum* KU-112 (abbreviated as St17), KU-120 (St24), and KU-122 (St25)^13^ using Br48 as a test isolate. Since *Rmg7* was located on the distal end of 2AL in which the *Pm4* locus resided, we assumed that *Rmg7* might be an allele of *Pm4*. PCR amplification and sequencing revealed that these three accessions shared a gene identical to *Pm4a*. In 93 F_2:3_ lines derived from St24 x Tat14 (*T. paleocolchicum* KU-156, susceptible to Br48), reactions to Br48 (conferred by *Rmg7*) perfectly co-segregated with the presence/absence of *Pm4a* (Fig. 2a) determined by KM200, another presence/absence PCR marker designed on *Rmg8-V2* (Fig. 1c, Extended Data Fig. 3).

To confirm that *Pm4a* recognize *AVR-Rmg8*, a protoplast cell death assay was performed. cDNA fragments of the two alternative splicing variants derived from *Pm4a* in St24 was inserted into pZH2Bik and established as pRmg7-V1 and pRmg7-V2, respectively. Fluorescence was not reduced in pAVR-Rmg8/pRmg7-V1 or pAVR-Rmg8/pRmg7-V2 combinations (Fig. 2b). When protoplasts were co-transfected with pAVR-Rmg8 and a mixture of pRmg7-V1 and pRmg7-V2, however, fluorescence was significantly reduced. This reduction was cancelled when pAVR-Rmg8 was replaced with pPWT3. These results suggest that *Pm4a* specifically recognizes *AVR-Rmg8*, and is therefore *Rmg7*.

### Distribution of *Rmg8* variants in common wheat

From here, we will call genes that can be amplified with KM171 or KM200 (including *Rmg8*, *Rmg7*, and *Pm4* alleles reported previously) as *Rmg8* variants collectively. As mentioned above, we previously screened 520 local landraces of common wheat by inoculation with Br48 and found 18 accessions that recognized *AVR-Rmg8*^16^. Although they assumed that these 18 accessions were *Rmg8* carriers, there remained a possibility that some of them might be *Rmg7* carriers because *Rmg7* also recognized *AVR-Rmg8*. In the present study we again screened a total of 526 local landraces (the 520 accessions plus 6 additional accessions) with Br48, and found 21 resistant accessions (the 18 accession plus 3 additional accessions). They were all susceptible to Br48ΔA8 but resistant to Br48ΔA8+eI, and therefore, confirmed to recognize *AVR-Rmg8* (Extended Data Table 1). Sequence analysis revealed that more than half of them (12 accessions) carried *Pm4f* as expected. However, the other accessions were composed of one *Pm4b* carrier and eight *Pm4a* carriers. These *Pm4a* carriers included three accessions (IL92, CP71, GR250) which had already been confirmed to carry a single resistance gene at the same locus as S-615, i.e. on 2BL^23^. To further check chromosomal locations of *Pm4a* in common wheat, we chose two *Pm4a* carriers (IL186, CP20) and crossed them with S-615. In the resulting F_2_ populations, resistant and susceptible seedlings segregated in 15:1 ratios (Extended Data Table 2), suggesting that they were carriers of *Rmg7* located on 2AL. Taken together, these results suggest that the chromosomal location of the *Pm4a* sequence is not restricted to 2AL; it resides on 2AL in some accessions but on 2BL in others.

To find other *Rmg8* variants, we screened the rest of the local landraces (505 susceptible accessions) with the PCR marker KM200, and found 7 accessions carrying *Pm4g* and 3 accessions carrying a new variant tentatively designated as *PM4_h1* (Extended Data Table 1). They were considered to be ineffective to MoT (Extended Data Table 1). *Pm4d* or *Pm4h* were not detected in our collection of common wheat local landraces.

These *Rmg8* variants were plotted on maps of Europe, Middle East, and Africa (Ethiopia). *Pm4f* and *Pm4a*, which are effective against MoT, were distributed from Middle East through southern Europe (Fig. 3a). On the other hand, *Pm4g*, which is ineffective against MoT, was distributed around mid-northern areas of Europe. The *Rmg8* variants were scarcely detected in accessions collected in Asia and the Americas (Extended Data Table 3).

**Fig. 3.**
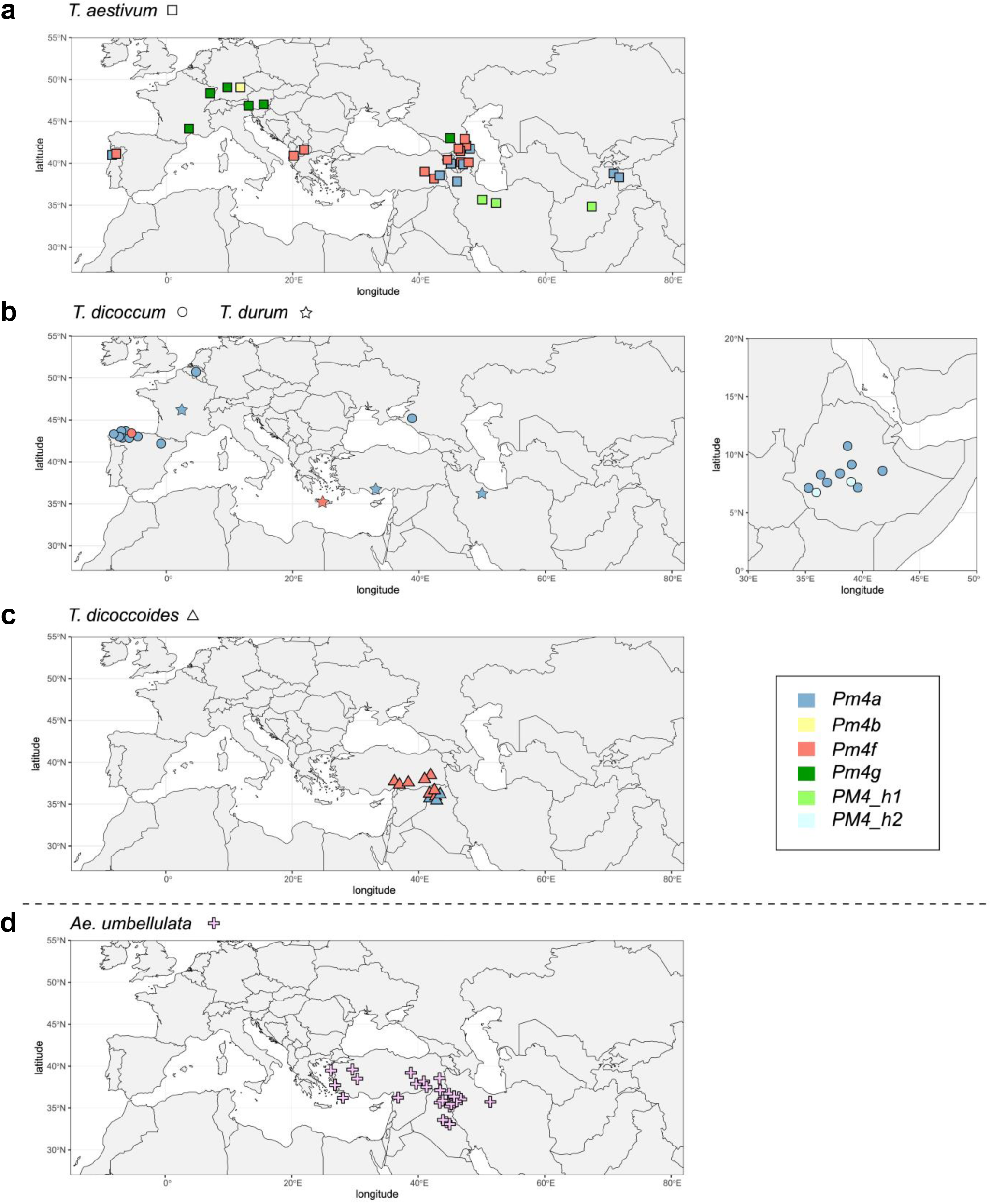
Geographical distribution of *Rmg8* variants. In common wheat (**a**), cultivated tetraploid wheat (**b**) and wild tetraploid wheat (**c**), accessions carrying *Pm4* “alleles” were plotted. The *Pm4* “alleles” were color-coded. In *Ae. umbellulata* (**d**), accessions recognizing *AVR-Rmg8* were plotted without differentiation of their *Rmg8* variants.

### Distribution of *Rmg8* variants in tetraploid wheat

To trace the origin of the *Rmg8* variants, we screened tetraploid wheat composed of 46 accessions of *T. dicoccoides*, 76 accessions of *T. dicoccum*, 72 accessions of *T. durum*, and 4 accessions of *T. paleocolchicum* with KM200. For accessions positive with KM200, the entire gene was amplified and sequenced. In the wild emmer wheat (*T. dicoccoides*) *Pm4f* was detected more frequently than *Pm4a* (Extended Data Table 3). In the cultivated emmer wheat (*T. dicoccum*), however, *Pm4a* extremely predominated over *Pm4f* (Extended Data Table 3). Their geographical distribution suggested that, after the domestication of emmer wheat, *Pm4a* was preferentially transmitted from Fertile Crescent to Spain and Ethiopia (Fig. 3b, c) probably due to its advantageous character – the resistance to wheat powdery mildew. A new variant designated tentatively as *PM4_h2* was found in two accessions of *T. dicoccum* collected in Ethiopia. *Pm4a* and *Pm4f* were also detected in *T. durum* (Extended Data Table 3).

**Distribution of *Rmg8* variants in *Aegilops* spp.**

To reveal the origin of the functional genes recognizing *AVR-Rmg8*, *Aegilops* spp. composed of 909 accessions were screened by inoculation. Accessions resistant to Br48 but weakly resistant or susceptible to Br48ΔA8 were determined to be carriers of functional *Rmg8* variants. Such accessions were found in *Ae. umbellulata*, *Ae. speltoides*, and *Ae. comosa* (Extended Data Tables 4, 5). It should be noted that, in *Ae. umbellulata*, the 27 accessions resistant to Br48 were either susceptible (26 accessions) or weakly resistant (1 accession) to Br48ΔA8, suggesting that they all recognize *AVR-Rmg8*. Geographically, they were distributed around Fertile Crescent and Turkey (Fig. 3d).

Six accessions were arbitrarily chosen from the 26 accessions mentioned above and crossed with susceptible accessions. In each F_2_ population resistant and susceptible seedlings segregated in a 3:1 ratio (Extended Data Table 6), suggesting that the resistance of each accession is controlled by a single major gene. In addition, crosses among resistant accessions yielded no susceptible F_2_ seedlings (Extended Data Table 6), which was consistent with an assumption that they were allelic at the same locus.

### Phylogenetic analysis of *Rmg8* variants

The resistance genes recognizing *AVR-Rmg8* (*Rmg8* homologs) were amplified from seven, two, and one accessions of *Ae. umbellulata*, *Ae. speltoides*, and *Ae. comosa,* respectively, and sequenced. *Rmg8* variants from these species were designated as *AeuRmg8*, *AesRmg8*, and *AecRmg8*, respectively. These nucleotide sequences were aligned with those of tetraploid and hexaploid wheat lines, and a phylogenetic tree was constructed using MEGA7 (Fig. 4a). SY-Mattis, a common wheat cultivar carrying *Pm4d*^18^, was included in the materials. *AeuRmg8* and *AecRmg8* were grouped into a cluster remote from the others while *AusRmg8* was clustered with those of *Triticum* spp. and formed a subcluster with *Pm4g*. This is reasonable because the S genome in *Ae. speltoides* is close to the B genome of *Triticum* spp.^24,25^. *Rmg8* variants in *Triticum* spp. except *Pm4g* formed another subcluster. *Pm4f* was located on the basal part of this large subcluster and composed of various haplotypes, suggesting that *Pm4f* emerged earlier than the others. The topology suggested that *Pm4a*, *Pm4d*, *PM4_h1*, and *PM4_h2* originated from *Pm4f*, and that *Pm4b* originated from *Pm4d*.

**Fig. 4.**
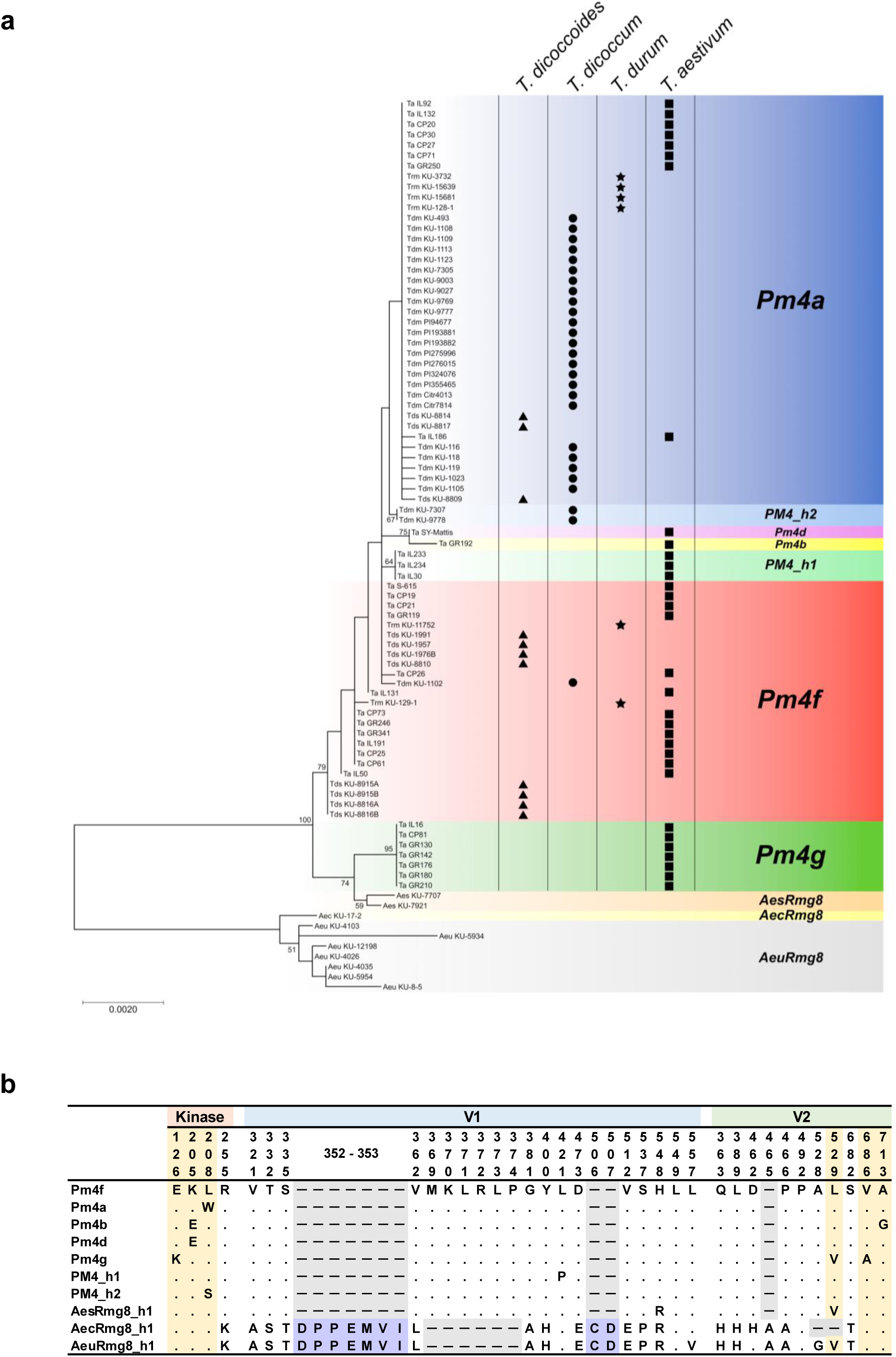
Phylogenetic relationships among *Rmg8* variants in *Triticum* and *Aegilops*. **a**, Maximum likelihood phylogenetic tree of *Rmg8* variants constructed using nucleotide sequences of the coding region. *Rmg8* variants in *Triticum* spp. are represented by their *Pm4* “allele” symbols. *PM4_h1* and *PM4_h2* are new variants which have not been reported previously. *AesRmg8*, *AecRmg8*, and *AeuRmg8* are *Rmg8* variants found in *Ae. speltoides*, *Ae. comosa*, and *Ae. umbellulata*, respectively. Bootstrap values (more than 50) from 1,000 replications are shown at nodes. Ta, *T. aestivum*; Trm, *T. durum*; Tdm, *T. dicoccum*; Tds, *T. dicoccoides*; Aes, *Ae. speltoides*; Aec, *Ae. comosa*; Aeu, *Ae. umbellulata*. **b**, Amino acid alignments of *Rmg8* variants detected in *Triticum* and *Aegilops* spp. AesRmg8_h1, AecAmg8_h1, and AeuRmg8_h1 are representatives of AesRmg8, AecAmg8, and AeuRmg8 detected in *Ae. speltoides* KU-7707, *Ae. comosa* KU-17-2, and *Ae. umbellulata* KU-4026, respectively.

Amino acid sequences of those variants are summarized in Fig. 4b with one representative from each of the *Aegilops* variants, i.e., AeuRmg8_h1 from *Ae. umbellulata* KU-4026, AesRmg8_h1 from *Ae. speltoides* KU-7707, and AecRmg8_h1 from *Ae. comosa* KU-17-2. The various haplotypes of *Pm4f* encoded the same protein with an identical amino acid sequence. Pm4a, Pm4d, PM4_h1, and PM4_h2 had a single amino acid substitution at different sites in comparison with Pm4f, suggesting that they emerged from Pm4f independently. Pm4b had two amino acid substitutions in comparison with Pm4f, but one of them was shared with Pm4d, supporting the idea that Pm4b evolved from Pm4d. AesRmg8_h1 was very similar to the Rmg8 variants in *Triticum* spp. while AeuRmg8_h1 and AecRmg8_h1 had large indels in comparison with them (Fig. 4b).

### Reactions of *Rmg8* variants to wheat blast and powdery mildew fungi

Reactions of representative *Rmg8* variants in *Triticum* spp. to wheat blast and wheat powdery mildew were tested using three MoT strains (Br48, Br48ΔA8, and Br48ΔA8+eI) and two Bgt isolates (Th1 and Th2) that were selected as representatives of 14 isolates collected in various locations in Japan (Extended Data Table 7). For *Pm4a*, common wheat cultivar Chancellor (Cc) and its near-isogenic line carrying *Pm4a* (Cc-Pm4a) were employed. Cc-Pm4a is a line (Khapli x Cc^8^) bred for mildew resistance by Briggle^26^. If our analysis mentioned above is correct, Cc-Pm4a should recognize *AVR-Rmg8*. Actually, Cc-Pm4a was resistant to Br48, susceptible to Br48ΔA8, and again resistant to Br48ΔA8+eI while Cc was susceptible to all of the three strains (Fig. 5). Against Bgt, Cc-Pm4a was resistant to Th2 while Cc was susceptible (Fig. 5), confirming that *Pm4a* is effective against Bgt. Cc-Pm4a was susceptible to another Bgt isolate Th1, suggesting that the avirulence gene corresponding to *Pm4a* is carried by Th2 but not by Th1. St24 is a tetraploid accession in which *Rmg7* (=*Pm4a*) was first identified^13^. Against the three MoT strains, St24 showed the same reactions as Cc-Pm4a as expected (Fig. 5). In addition, St24 showed strong resistance to the Bgt isolates (Fig. 5). Taken together, we confirmed that *Pm4a* is effective to both MoT and Bgt.

**Fig. 5.**
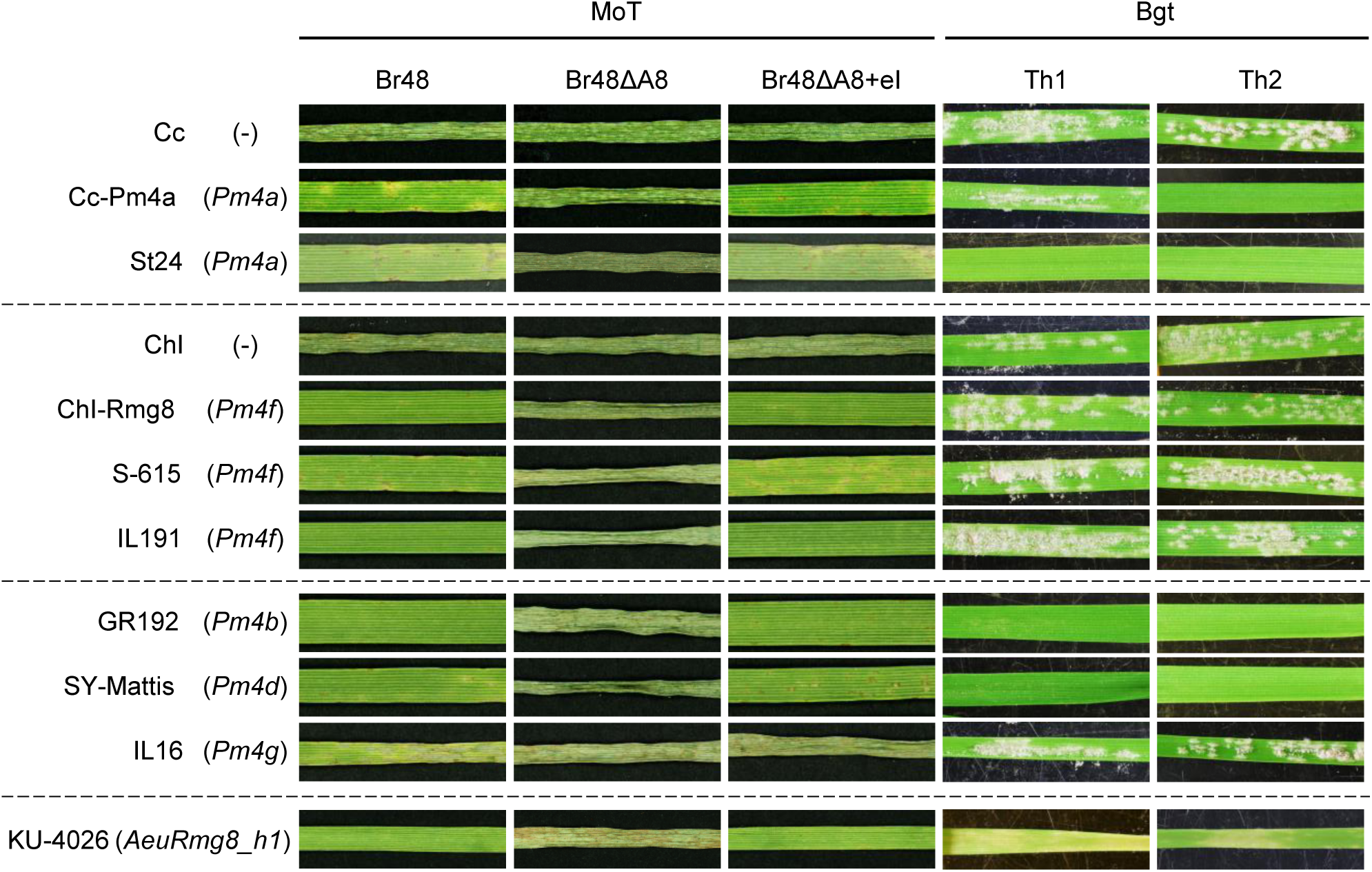
Reactions of *Rmg8* variants to wheat blast (MoT) and wheat powdery mildew (Bgt) fungi. Wheat accessions carrying *Rmg8* variants were inoculated with Br48 (wild MoT isolate), Br48ΔA8 (disruptant of *AVR-Rmg8*), and Br48ΔA8+eI (transformant of Br48ΔA8 carrying re-introduced *AVR-Rmg8*), and incubated for five days, or were inoculated with Th2 (Bgt carrying *AvrPm4a,* the putative avirulence gene corresponding to *Pm4a*) and Th1 (Bgt carrying *avrPm4a*, the non-functional allele of *AvrPm4a*), and incubated for eight days. *Rmg8* variants carried by the wheat lines are shown in parentheses. St24 and KU-4026 are a tetraploid accession and an *Ae. umbellulata* accession, respectively, and the others are common wheat lines. Cc-Pm4a is a near-isogenic line of cv. Chancellor (Cc) carrying *Pm4a* while ChI-Rmg8 is a near-isogenic line of cv. Chikugoizumi (ChI) carrying *Rmg8*.

For *Pm4f,* common wheat cultivar Chikugoizumi (ChI) and its near-isogenic line carrying *Rmg8* (=*Pm4f*) (ChI-Rmg8)^27^ were employed. ChI-Rmg8 was resistant to Br48, susceptible to Br48ΔA8, and again resistant to Br48ΔA8+eI as expected while ChI was susceptible to all of the three strains (Fig. 5). Other *Pm4f* carriers (S-615 and IL191) showed the same reactions. By contrast, these *Pm4f* carriers were all susceptible to Th1 and Th2 (Fig. 5). Furthermore, ChI-Rmg8 were susceptible to all Japanese Bgt isolates tested (Extended Data Table 7). Taken together, we concluded that *Pm4f* is effective to MoT but ineffective to Bgt.

GR192 carrying *Pm4b* and SY-Mattis carrying *Pm4d* were resistant to Br48, susceptible to Br48ΔA8, and resistant to Br48ΔA8+eI (Fig. 5), indicating that these alleles recognize *AVR-Rmg8*. They were also resistant to the Bgt isolates (Fig. 5). These results suggest that *Pm4b* and *Pm4d* are effective to both MoT and Bgt. On the other hand, IL16 carrying *Pm4g* was susceptible to all of the MoT strains and Bgt isolates tested (Fig. 5, Extended Data Table 7), suggesting that *Pm4g* is ineffective to both MoT and Bgt.

*Ae. umbellulata* KU-4026 carrying *AeuRmg8_h1* was resistant to Br48, susceptible to Br48ΔA8, and resistant to Br48ΔA8+eI (Fig. 5), confirming that *AeuRmg8_h1* recognizes *AVR-Rmg8*. When inoculated with the Th1 and Th2, primary leaves of KU-4026 became slightly chlorotic, but produced conidia enough to proceed to the next infection cycle (Fig. 5). KU-4026 showed similar reactions to all Japanese Bgt isolates tested (Extended Data Table 7). These results suggest that *AeuRmg8_h1* is effective to MoT but ineffective to Bgt.

## Discussion

In the present study we isolated *Rmg8*, the only genetic factor that has been identified as a major gene for resistance to MoT and supposed to be effective against wheat blast in farmer’s fields^27^. Intriguingly, *Rmg8* was identical to *Pm4f*, which was reported to be an “allele” of *Pm4*, a gene for resistance to wheat powdery mildew^18^. *Rmg8* was located on 2BL^14,27^ (Fig. 2a) while *Pm4a*, *Pm4b*, and *Pm4d* were reported to reside on 2AL^21,22^. This apparent discrepancy could be explained by considering that *Pm4f* was not a genetically identified allele but was found through PCR amplification and sequencing. We suggest that *Rmg8* is a homoeologous gene of *Pm4* alleles on 2AL. We further isolated *Rmg7* located on 2AL and found that this gene is identical to *Pm4a*. This is reasonable because *Rmg7* and *Rmg8* have been inferred to be homoeologous genes^14^. Intriguingly, the *Pm4a* sequence was also detected at the *Rmg8* locus on 2BL in some accessions (Extended Data Table 1). The *Pm4a* gene in these accessions should be considered to be *Rmg8* from the viewpoint of Mendelian genetics, but is identical to *Rmg7* at the molecular level.

The *Pm4* “alleles” tested were divided into three groups from the viewpoint of reactions to MoT and Bgt. The first one composed of *Pm4a*, *Pm4b*, and *Pm4d* was effective against both MoT and Bgt while the second one, *Pm4f*, was effective to MoT but ineffective to Bgt (Fig. 5). The third one, *Pm4g*, was ineffective to both MoT and Bgt (Fig. 5). *Pm4a*, *Pm4b*, and *Pm4d* have been identified as genes for resistance to Bgt and used for breeding. On the other hand, *Pm4f* and *Pm4g* were suggested to be susceptible “alleles” against Bgt^18^. In addition, carriers of these “alleles” were susceptible to all Japanese Bgt isolates tested (Extended Data Table 7). One hypothesis to explain this general susceptibility would be that they had been effective against Bgt at the time of their emergence, but were later overcome by newly evolved virulent races. However, this scenario implies that their corresponding avirulence genes have been eliminated from the Bgt populations in both Europe and Far East, and therefore requires a wide cultivation of wheat lines carrying these ‘resistance genes’. Considering their low frequencies in local landraces and no record of wide cultivation of such cultivars, however, such perfect elimination is unlikely to occur. Therefore, *Pm4f* and *Pm4g* are considered to have been ineffective against Bgt from the time of their emergence. The phylogenetic tree (Fig. 4) suggested that *Pm4a*, *Pm4b*, and *Pm4d* evolved from *Pm4f*. Taken together with the above discussion, we suggest that these *Pm4* alleles for resistance to powdery mildew have evolved from *Pm4f* through gaining an ability to recognize Bgt. The predominated distribution of *Pm4a* over *Pm4f* in the cultivated emmer wheat in contrast to their distribution in the wild emmer wheat (Figs. 3, 4) may be attributable to preferential transmission of *Pm4a* carriers by peoples who noticed the advantage of powdery mildew resistance conferred by this allele.

The gain of the ability to recognize Bgt was caused by a single amino acid substitution (Fig. 4b), and resulted in the generation of the alleles expressing resistance to both MoT and Bgt (Fig. 5). There are two additional examples suggesting close associations of recognition of *P. oryzae* and *B. graminis*. Two amino acid deletion of *Rwt4*, a gene for resistance to an *Avena* isolate of *P. oryzae*, resulted in a gain of resistance to Bgt^28,29^. *Mla3*, an allele at the *Mla* locus conditioning the resistance of barley to *B. graminis* f. sp. *hordei* (Bgh, the barley powdery mildew fungus), recognized *PWL2*, an avirulence gene derived from *P. oryzae* pathotype *Oryza* (the rice blast fungus)^30^. Mechanisms of such dual specificity with *P. oryzae* and *B. graminis* should be elucidated at the level of molecular structures.

Functional *Rmg8* variants were also detected in *Ae. umbellulata* (U genome), *Ae. speltoides* (S genome), and *Ae. comosa* (M genome). This result suggests that the prototype of *Rmg8* equipped with the function for recognizing *AVR-Rmg8* was established before the differentiation of the A, B, U, S, and M genomes in the *Triticum* – *Aegilops* complex, which was estimated to be 5–6 million years ago^31^. This implies that this gene has maintained the function for recognizing *AVR-Rmg8* for 5–6 million years without infection pressure exerted by MoT because MoT first emerged in 1985. However, it seems unlikely that a resistance gene has maintained its function for such a long time under no infection pressure by pathogens. One possibility is that *Rmg8* and its variants had been interacting with *P. oryzae* before the differentiation into pathotypes, and after the differentiation, have been interacting with pathotype(s) that maintained *AVR-Rmg8*. The most probable candidate of such pathotypes is inferred to be MoL (*Lolium* pathotype) with three reasons. First, MoL is phylogenetically the closest to MoT^32^. Second, functional *AVR-Rmg8* is widely distributed in the population of MoL^33^. Third, its hosts (Italian ryegrass and perennial ryegrass) are widely distributed in Middle East and southern Europe^34–36^ where functional *Rmg8* variants are frequently found (Fig. 3). Another possibility is that the *Rmg8* variants have been effective against pathogens other than the blast fungus (and the powdery mildew fungus). It should be noted that *Sr33* in wheat and *Sr50* in rye, genes for resistance to stem rust, are homologs of *Mla*, a barley gene for resistance to Bgh^37,38^. Also, an allele at the *Mla* locus, *Mla8*, was shown to be effective against wheat stripe rust^39^.

In Introduction we raised a question why common wheat accessions in Europe and Middle East have maintained *Rmg8*, a gene for resistance to MoT. The present study revealed that “*Rmg8*” detected in those accessions was composed of *Pm4a*, *Pm4b*, and *Pm4f*. We suggest that about a half of them (carriers of *Pm4a* and *Pm4b*) have maintained these genes due to their effectiveness against wheat powdery mildew. The maintenance of *Pm4f* in the other accessions may be explained by the same reasoning as the *Rmg8* variants in *Aegilops* spp; *Pm4f* may have been effective against MoL or other pathogens. It is suggestive that *Pm4f* is distributed in similar regions as the *Rmg8* variants in *Aegilops* spp., i.e., warm areas around the same latitude (Fig. 3).

The evolutionary process of *Rmg8* inferred from the present study is summarized in Fig. 6. The prototype of *Rmg8* gained an ability to recognize *AVR-Rmg8* before the differentiation of *Triticum* and *Aegilops*. It then differentiated into variants including *Pm4g* and *Pm4f*. Some variants derived from *Pm4f* gained an ability to recognize Bgt, and evolved into *Pm4a*, *Pm4d*, and *Pm4b*, genes for resistance to wheat powdery mildew. Finally, when MoT emerged in 1985, those *Rmg8* variants appeared as genes for resistance to wheat blast because they recognized an effector encoded by *AVR-Rmg8* of MoT. This figure illustrates an evolutionary process in which a resistance gene has expanded its target pathogens. The present study also provides perspectives from the viewpoint of breeding. When a resistance gene to a known pathogen is cloned, nucleotide sequences of its “susceptible alleles” should be also clarified. If a “susceptible allele” maintaining an ORF is distributed in the crop population in a certain frequency, it may be a functional resistance gene against other pathogen(s) which is prevailing now or those which will emerge in future. Conversely, when a new disease emerges, resistance genes against its causal agent (a new pathogen) may be found among known resistance genes against currently prevailing pathogens.

**Fig. 6.**
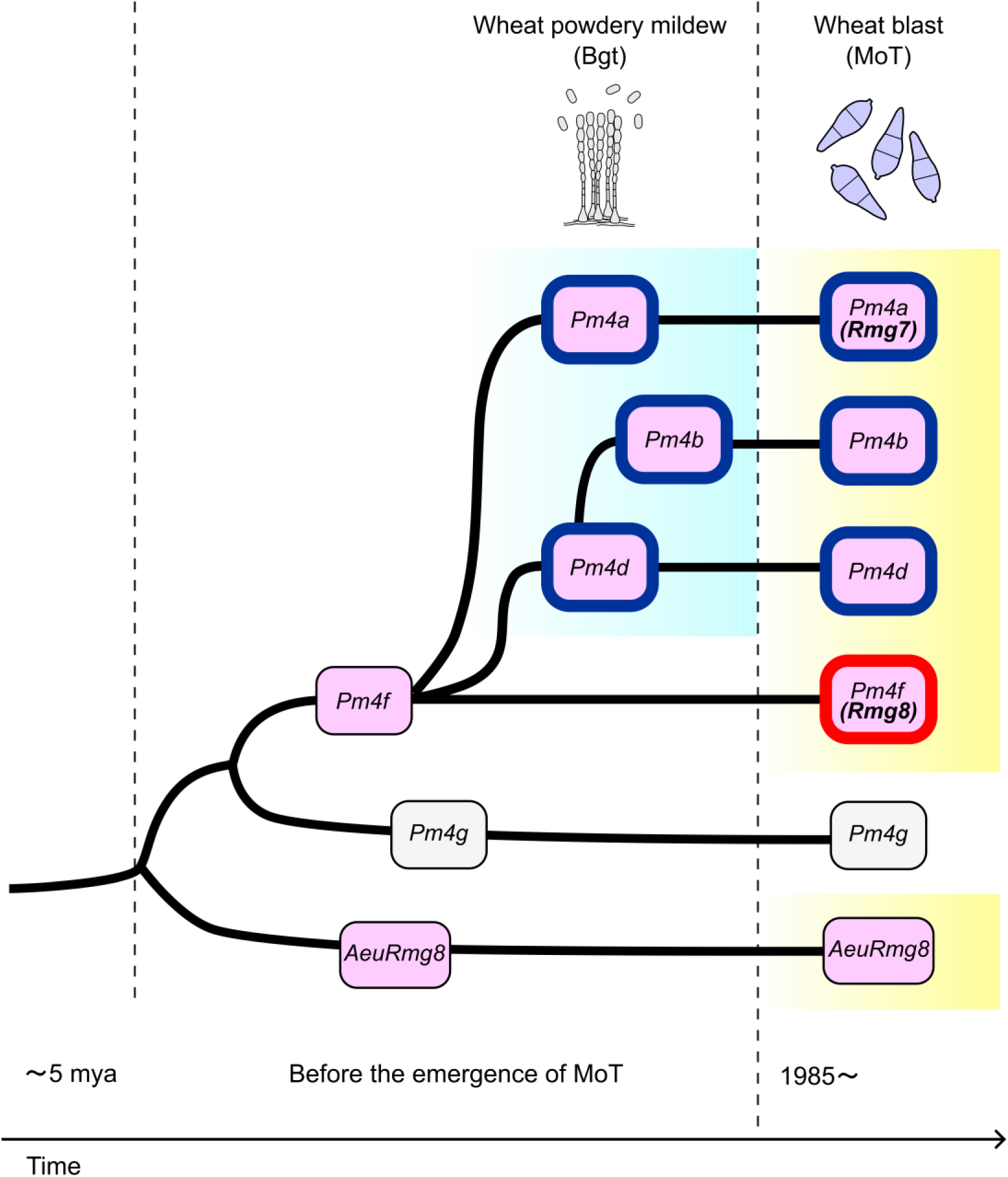
A model illustrating evolutionary processes in which resistance gene *Rmg8* has gained new target pathogens through differentiation of variants. The *Rmg8* variants painted in pink and enclosed in a blue rectangle indicate those with functions for recognizing wheat blast (MoT) and wheat powdery mildew (Bgt), respectively. The red rectangle indicates recognition as a useful gene for resistance to wheat blast. mya, million years ago.

## Methods

### Plant materials

Parental cultivars for mapping of *Rmg8*, *Triticum aestivum* cv. S-615 and cv. Shin-chunaga (Sch), were provided by K. Tsunewaki, Emeritus professor at Kyoto University, Japan. Parental accessions for mapping of *Rmg7*, *T. dicoccum* St24 (accession No. KU-120) and *T. paleocolchicum* Tat14 (KU-156), were provided by S. Sakamoto, Emeritus professor at Kyoto University. *Hordeum vulgare* cv. Golden Promise (GP) for protoplast assay and *T. aestivum* cv. Fielder (KT020-061) for transformation were provided by K. Sato, Okayama University, Japan, and the National BioResource Project –Wheat (NBRP) (https://shigen.nig.ac.jp/wheat/komugi/), Japan, respectively. *T. aestivum* cv. Chancellor (Cc) and its near isogenic line Cc-Pm4a carrying *Pm4a* (=Khapli x Cc^8^ produced by Briggle^26^) were provided by U. Hiura, Emeritus professor at Okayama University. *T. aestivum* cv. Chikugoizumi (ChI) and its near-isogenic line carrying *Rmg8* (ChI-Rmg8)^27^ were produced in the BRAIN project (see Acknowledgments), and maintained at NARO (National Agriculture and Food Research Organization), Japan. *T. aestivum* cv. SY-Mattis, one of the accessions analyzed by the wheat pangenome project^40^, was provided by John Innes Centre to S. Nasuda and maintained at Kyoto University. The 526 local landraces of *T. aestivum* used for the distribution analysis were a collection of K. Kato, Okayama University, Japan. Original providers of the *T. aestivum* accessions carrying the *Rmg8* variants are shown in Extended Data Table 1. The accessions of tetraploid wheat used for the distribution analysis were collections of N. Mori, Kobe University, and S. Nasuda, Kyoto University. The tetraploid accessions carrying the *Rmg8* variants are shown in Fig. 4. Among them accessions with the prefix KU-were provided by NBRP while those with the prefixes PI and Citr were provided by the U.S. National Plant Germplasm System. The 909 accessions of *Aegilops* spp. used for screening for functional *Rmg8* variants were provided by NBRP.

### Fungal materials

Wheat blast strains used for infection assay were *Pyricularia oryzae* pathotype *Triticum* wild isolate Br48 collected in 1990 in Brazil, Br48ΔA8_d6 (abbreviated as Br48ΔA8), a disruptant of *AVR-Rmg8* derived from Br48^16^, and Br48ΔA8+eI-3 (abbreviated as Br48ΔA8+eI), a transformant of Br48ΔA8 carrying the eI type of *AVR-Rmg8*^41^. They have been maintained on sterilized barley seeds at Kobe University.

Wheat powdery mildew strains used were wild isolates of *Blumeria graminis* f. sp. *tritici* collected in various regions in Japan (Extended Data Table 7). They were purified through single-conidium isolation and have been maintained at 4°C on primary leaves of *T. aestivum* cv. Norin 4 through subculturing.

### Inoculation with wheat blast strains

Seeds of *Triticum* and *Aegilops* spp. were pregerminated on a moistened filter paper for 24h. Germinated seeds of *Triticum* spp. were sown in vermiculite supplied with liquid fertilizer in a seedling case (5.5 x 15 x 10 cm) and grown at 22℃ with a 12-h photoperiod of fluorescent lighting for 8 days. Germinated seeds of *Aegilops* spp. accessions were sown in the seedling case filled with Sakata Prime Mix soil (Sakata, Japan) and grown at 22℃ with a 12-h photoperiod of fluorescent lighting for 21 days. Primary leaves of eight-day-old wheat seedlings or first to third leaves of 21-day-old *Aegilops* seedlings were fixed onto a plastic board with rubber bands just before inoculation. Conidial suspensions (1×10^5^ conidia/ml) prepared as described previously^13^ were sprayed onto fixed leaves using an air compressor. The inoculated seedlings were incubated in a sealed box under dark and humid conditions at 22°C for 24h, then transferred to dry conditions with a 12h photoperiod of fluorescent lighting, and incubated for additional 3-5 days at 22°C. Four to six days after inoculation, symptoms were evaluated based on the color of lesions and the affected leaf area. The affected area was rated by six progressive grades from 0 to 5: 0 = no visible evidence of infection; 1 = pinhead spots; 2 = small lesions (<1.5 mm); 3 = scattered lesions of intermediate size (<3 mm); 4 = large typical blast lesions; and 5 = complete blighting of leaf blades. A disease score (infection type) was designated by combining a number which denotes the size of lesions and a letter or letters indicating the lesion color, i.e., ‘B’ for brown and ‘G’ for green. Infection types 0 to 5 with brown lesions were considered to be resistant while infection types 3G, 4G, and 5G were considered to be susceptible. Infection type 3BG accompanies by a mixture of brown and green lesions were taken as weakly resistant.

### Inoculation with powdery mildew isolates

Seeds of test plants were sown in autoclaved soil in 2×30cm or 2×35 cm test tubes. Eight days after sowing, primary leaves were inoculated with conidia from eight-day-old colonies using writing brushes. The seedlings were incubated at 22 °C in a controlled-environment room with a 12-h photoperiod of fluorescent lighting. Seven to eight days after inoculation, infection types were recorded using five progressive grades from 0 to 4: 0, no mycelial growth or sporulation; 1, scant sporulation; 2, reduced sporulation; 3, slightly reduced sporulation; 4, heavy sporulation.

### Mapping of *Rmg8* and *Rmg7*

A total of 165 F_2:3_ lines derived from a cross between S-615 and Sch were used for mapping of *Rmg8*. Twenty seeds were retrieved from each F_2:3_ line and subjected to infection assay with Br48 for phenotyping. Another set of 20 seeds was retrieved from each F_2:3_ line, sown in vermiculite, and grown at 22℃ for 7 days. Seven-day-old primary leaves were bulked, and subjected to DNA extraction by the CTAB method. For detecting polymorphisms between S-615 (*Rmg8*) and Sch (*rmg8*), total RNA was extracted from their primary leaves using Maxwell RSC Plant RNA Kit (Promega). Sequence libraries were generated by NEBNext Ultra II Directional RNA Library Prep Kit, and sequenced using Illumina Hiseq (paired-end) by sequencing service of Novogene, Japan. Sequence reads of S-615 and Sch were aligned to the reference genome of Chinese Spring version 1.1 and 2.0^42,43^ using HISAT2 (v2.1.1), and variants were called by samtools (v1.8) to generate VCF files. Using the VCF files, Cleaved Amplified Polymorphic Sequence (CAPS) and presence/absence markers were developed. Marker fragments were amplified from genomic DNA of the parental cultivars and the F_2:3_ lines using 2x Quick Taq HS DyeMix (TOYOBO, Osaka, Japan) following the manufacturer’s instructions. Fragments amplified with primers for CAPS markers were digested with appropriate restriction enzymes supplied by Takara Bio (Kusatsu, Japan) or New England Biolabs Japan (Tokyo, Japan) (Extended Data Table 8). PCR products were electrophoresed in 0.7-2.0% agarose gels and stained with ethidium bromide for visualization. MAPMAKER/EXP version 3.0 was used for constructing a genetic map^44^. The logarithm-of-odds (LOD) threshold for declaration of linkage was set at 4.0. Genetic distance was calculated with the Kosambi function.

For mapping of *Rmg7*, RNA sequencing of St24 and Tat14 was performed in a similar way as mentioned above. Sequenced reads were aligned to the reference genome of *Triticum dicoccoides* cv. Svevo (EnsemblPlants, https://plants.ensembl.org/Triticum_turgidum /Info/Index) to develop genetic markers.

### Screening for a transcript derived from *Rmg8* based on an association analysis with susceptible F_2:3_ lines

To detect candidate genes for *Rmg8*, we carried out association analysis of expressed genes. First, cDNA sequence of S-615 transcripts was generated by sequencing service (GeneBay, Japan). Base calling of ONT reads was performed on FAST5 files using Guppy (Oxford Nanopore Technologies). Hybrid de novo assembly was performed by rnaSPEdes^45^ using ONT reads and Illumina short reads both, resulting in 161,852 transcripts. ORFs coding more than 300 amino acids in these transcripts were predicted by TransDecodar^46^ and cd-hit^47^, resulting in a reference cDNA sequence set of S-615 composed of 27,205 transcripts.

Next, we chose nine F_2:3_ lines susceptible to Br48 arbitrarily, extracted total RNA from three leaves of each F_3_ line using Maxwell RSC Plant RNA Kit (Promega), prepared sequence libraries, and sequenced them using Illumina Hiseq4000 (150 bp Paired-End reads) in a similar way as mentioned above. The presence/absence analysis was carried out based on the transcripts per million (TPM) value of the transcripts for selecting genes which were expressed in S-615 but not in either Sch or the nine susceptible F_2:3_ lines.

### Cell death assay with barley protoplasts

The genomic sequence of *Rmg8* (*Rmg8-genome*) and the transcript variants of *Rmg8* and *Rmg7* (Rmg8-V1, Rmg8-V2, Rmg7-V1 and Rmg7-V2) were employed for protoplast cell death assays. The fragment of *Rmg8-genome* was amplified from genomic DNA of S-615. Rmg8-V1 and Rmg8-V2 were amplified from cDNA of S-615 while Rmg7-V1 and Rmg7-V2 were amplified from cDNA of St24. RNA extraction and cDNA synthesis were performed as mentioned in the previous sections. The ORFs of *PWT3* and *AVR-Rmg8* without signal peptides were amplified from genomic DNA of Br58^5^ and Br48, respectively. Primers used for these PCR reactions are shown in Extended Data Table 8. All of these fragments were cloned into the *Kpn*I site in the pZH2Bik vector using In-Fusion cloning (Takara) so as to be driven by the rice ubiquitin promoter, resulting in pRmg8-genome, pRmg8-V1, pRmg8-V2, pRmg7-V1, pRmg7-V2, pPWT3, and pAVR-Rmg8. Established plasmids were extracted by NucleoBond Xtra Maxi (Macherey-Nagel, Düren, Germany). Barley cultivar GP was employed as a recipient of transgenes because barley epidermis could be peeled off more easily than wheat epidermis for releasing protoplasts. Mesophyll protoplasts were prepared from eight-day-old primary leaves of GP. Transfection assays with these plasmids were performed as described in Saur et al.^19^. Briefly, plasmids containing AVR and resistance genes were introduced into the GP protoplasts with a plasmid containing the luciferase gene (pAHC17-LUC) via the polyethylene glycol treatment. After 18 hours incubation at 20°C in the dark, the protoplasts were lysed, and luciferase activity in the resulting cell extracts was measured for 1 second/well on the Tristar 3 luminometer mode (Berthold). The measured luminescence was normalized using the negative control in which the AVR gene was substituted with the empty pZH2Bik vector. This experiment was repeated four times independently.

### Production of transgenic plants

pRmg8-V1, pRmg8-V2, and pRmg8-genome were introduced into *T. aestivum* cv. Fielder via the Agrobacterium*-*mediated transformation as described by Ishida et al.^48^ Insertions of transgenes were checked by PCR with the HPT primers (Extended Data Table 8). We obtained three, eleven, and three T_1_ lines carrying *Rmg8-V1*, *Rmg8-V2*, and *Rmg8-genome*, respectively. Transgenic T_1_ seedlings were inoculated with Br48, Br48ΔA8, and Br48ΔA8+eI to evaluate functions of transgenes.

### Sequencing of *Rmg8* variants and phylogenetic analysis

In the distribution analyses in *Triticum* spp. all test accessions were screened with KM171 and KM200, and those with amplicons were subjected to sequence analyses irrespective of their phenotypes (resistant or susceptible). In the analysis of *Aegilops* spp. all accessions were first screened by inoculation with Br48 and Br48ΔA8, and those recognizing *AVR-Rmg8* were subjected to sequence analyses. Two primer pairs were used to amplify two different regions of the *Rmg8* genomic fragment, one encoding exons 1 to 5 and the other encoding exons 6 and 7. These fragments were inserted into the *EcoR*V site of pBSIISK+, sequenced with ABI capillary sequencers, and aligned with MAFFT (v7.520). Coding sequences were extracted from obtained sequences and concatenated. A maximum likelihood tree was constructed using MEGA X^49^ with 1,000 bootstrap replicates. Primers used in this section are listed in Extended Data Table 8.

## Data availability

Sequence data for the genes described in the present study can be found in the GenBank/EMBL database under the accession numbers LC779671, LC779672, LC779673, and LC779674. All plasmids, plant lines, and fungal strains generated in this work are available from the authors upon request.

## Acknowledgments

We thank Izumi Chuma (Obihiro University, Japan), Kaori Nakajima (Mie Prefecture Agricultural Research Institute, Japan), Atsushi Ohta (Kyoto University, Japan), Kaichi Uchihashi (Hyogo Prefectural Technology Center for Agriculture, Japan), Hisashi Tsujimoto (Tottori University, Japan), and Tomomori Kataoka (National Agricultural Research Center for Kyushu Okinawa Region, Japan) for providing powdery-mildewed wheat leaves collected in fields. We also thank Paul Nicholson (John Innes Centre, U.K.), Barbara Valent (Kansas State University, U.S.A.), and Brian Staskawicz (University of California, Berkeley, U.S.A.) for valuable suggestions on the manuscript. *Aegilops* spp. accessions were provided by the National BioResource Project–Wheat with support in part by the National BioResource Project of the MEXT, Japan. Computations were partially performed on the NIG supercomputer owned by National Institute of Genetics, Research Organization of Information and Systems. This research was supported by the research program on development of innovative technology grants (JPJ007097) from the project of the Bio-oriented Technology Research Advancement Institution (BRAIN) and a grant, “International collaborative research project for solving global issues”, from Agriculture, Forestry and Fisheries Research Council Secretariat, Ministry of Agriculture, Forestry and Fisheries (MAFF), Japan.

