## Supplemental data for "Evolution of wheat blast resistance gene *Rmg8* accompanied by differentiation of variants recognizing the powdery mildew fungus"

**a**

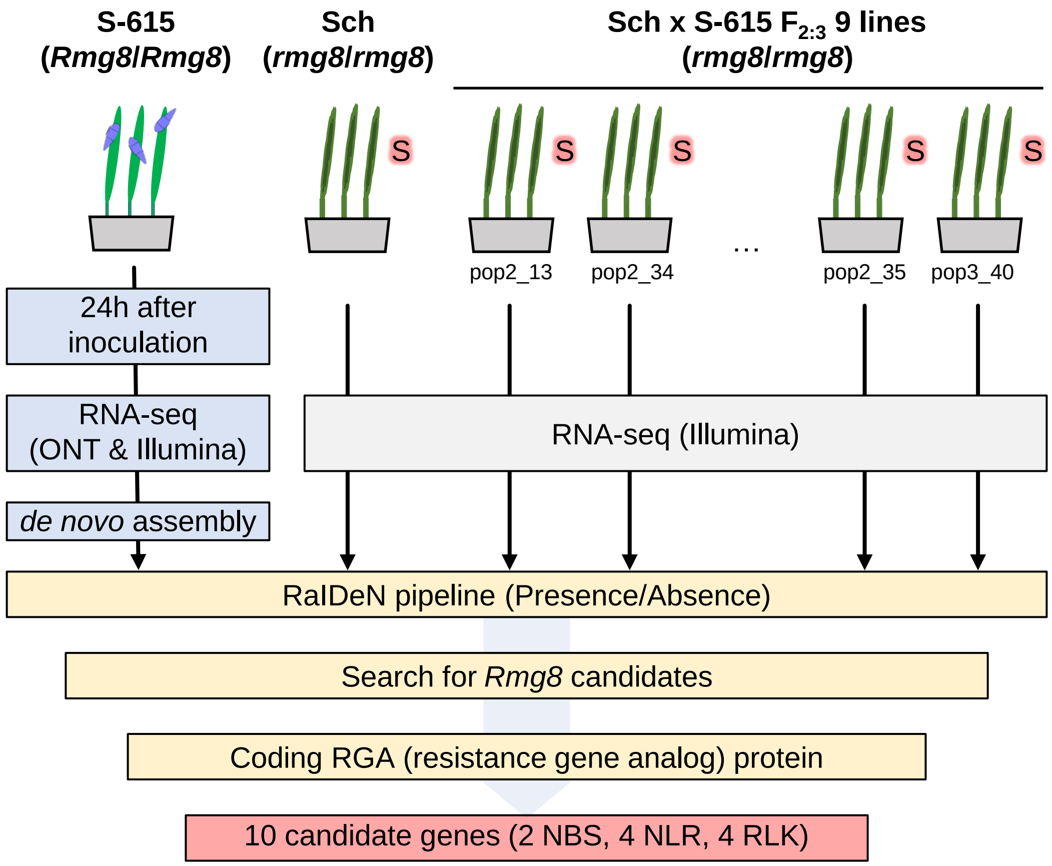


**b**

| ID | ORF name | Domain prediction | Peptide homolog in *T. turgidum* cv. Svevo | Chromosome | Mapping with Sch × S-615 | | Association |
|---|---|---|---|---|---|---|---|
| | | | | | Marker | co-segregation with *Rmg8* | |
| Can-A | NODE_34172 | NBS | TRITD2Bv1G265560 | 2B | KM30, KM109 | Yes | No |
| Can-B | NODE_47471 | NBS | TRITD2Bv1G264480 | 2B | KM27 | Yes | No |
| Can-C | NODE_2794 | NLR | TRITD2Bv1G265560 | 2B | KM30, KM109 | Yes | No |
| Can-D | NODE_43629 | NLR | TRITD2Bv1G264480 | 2B | KM27 | Yes | No |
| Can-E | NODE_44374 | NLR | TRITD6Bv1G223090 | 6B | - | - | - |
| Can-F | NODE_13794 | NLR | TRITD2Bv1G264470 | 2B | KM138, KM140 | Yes | No |
| Can-G | NODE_6206 | RLK | TRITD4Av1G219830 | 4A | - | - | - |
| Can-H | NODE_12802 | RLK | TRITD3Av1G028450 | 3A | - | - | - |
| **Can-I** | **NODE_16006** | **RLK** | **TRITD2Bv1G265720** | **2B** | **KM171** | **Yes** | **Yes** |
| Can-J | NODE_8088 | RLK | TRITD3Av1G247380 | 3A | - | - | - |

**Extended Data Fig. 1. Search for *Rmg8* candidate genes through association analyses of expressed genes with susceptible $F_{2:3}$ lines. a**, An outline of screening of candidate genes. **b**, A list of *Rmg8* candidate genes. NBS, Nucleotide-binding domain; NLR, Nucleotide-binding domain and leucine-rich repeat; RLK, Receptor-like kinase.

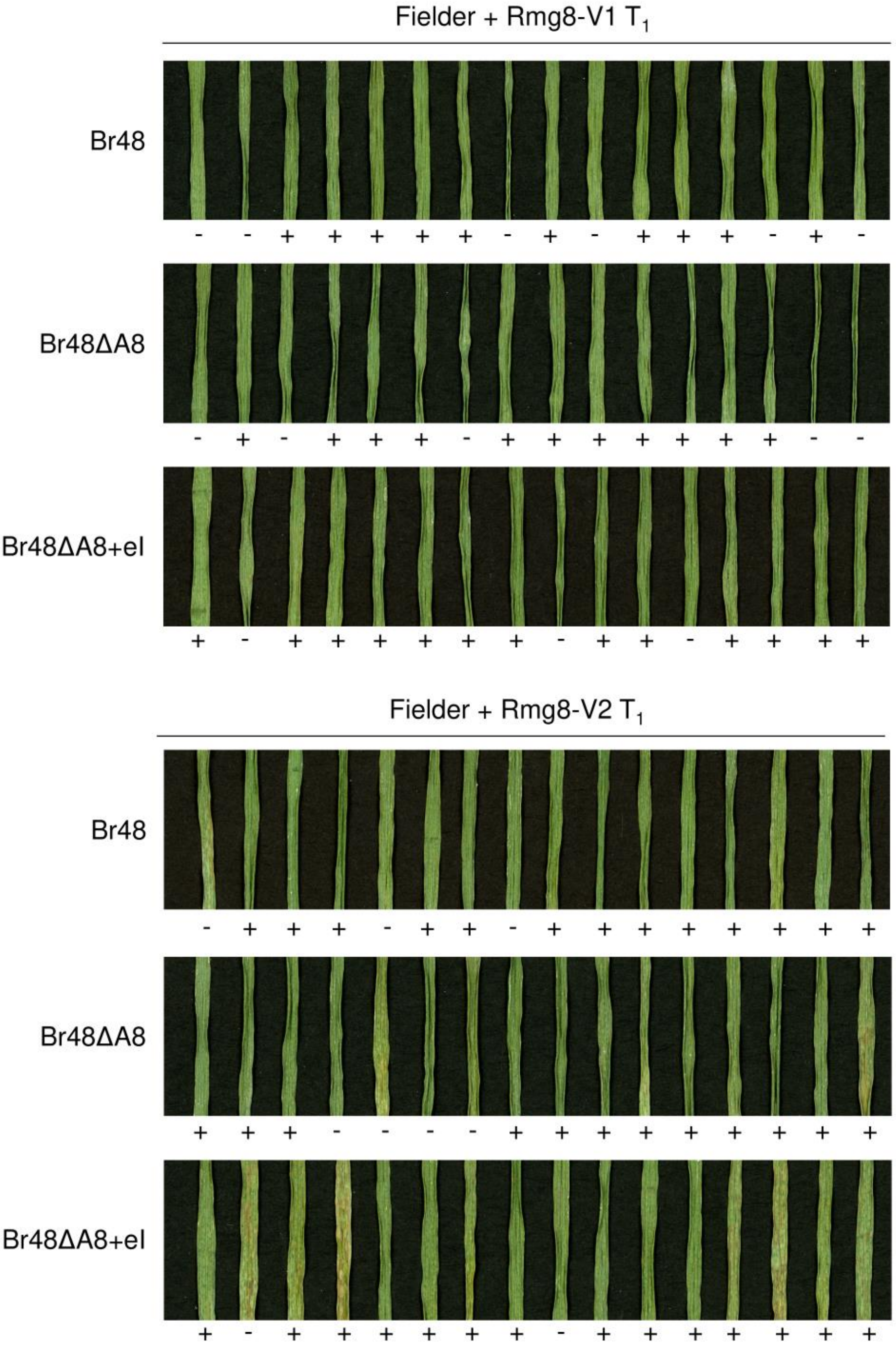


**Extended Data Fig. 2. Reactions of $T_1$ transformants carrying *Rmg8-V1* and *Rmg8-V2*.** $T_1$ individuals derived from transformation of Fielder with the *Rmg8-V1* CDS (Fielder+Rmg8-V1) or with *Rmg8-V2* CDS (Fielder+Rmg8-V2) were inoculated with Br48, Br48ΔA8, and Br48ΔA8+eI, and incubated for five days. Presence (+)/absence (-) of the transgene confirmed by PCR with the HPT primers are shown below the panels.

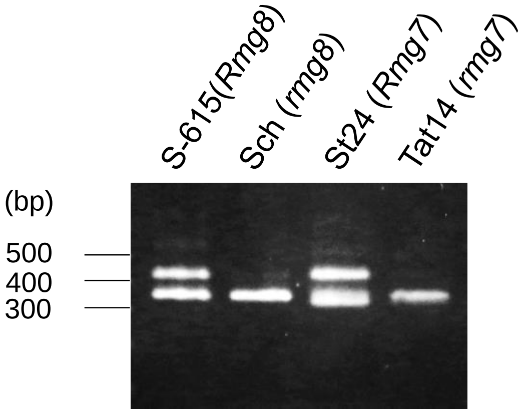


**Extended Data Fig. 3. PCR products amplified with KM200 primers.** Genomic DNAs of common wheat (S-615, Sch) and tetraploid wheat (St24, Tat14) were subjected to amplification with KM200 primers, and resulting amplicons were run on a 2% gel for 30 min. The 424bp fragment is amplified from both of the *Rmg8* carrier and the *Rmg7* carrier, but not from the noncarriers. The 350bp fragment is amplified from all cultivars/accessions irrespective of their genotypes, and can be used as an indicator of successful PCR reactions.

Extended Data Table 1. *Rmg8* variants detected in local landraces of common wheat

| Accession[a] | Origin | Donor[b] | Accession No. | *Rmg8* variant | Infection type[c] with Br48 | Infection type[c] with Br48ΔA8 | Infection type[c] with Br48ΔA8+eI | Recognition of *AVR-Rmg8* |
|---|---|---|---|---|---|---|---|---|
| GR119* | Albania | JIC | W7880 | *Pm4f* | 0 | 2B | 0 | + |
| IL191* | Armenia | NBRP | KU-1649 | *Pm4f* | 0 | 5G | 0 | + |
| CP19* | Azerbaijan | VIR | WIR-39310 | *Pm4f* | 1B | 5G | 0 | + |
| CP21* | Azerbaijan | VIR | WIR-39844 | *Pm4f* | 0-1B | 5G | 0-1B | + |
| CP25 | Azerbaijan | VIR | WIR-39934 | *Pm4f* | 1B | 5G | 0-1B | + |
| CP26* | Azerbaijan | VIR | WIR-40184 | *Pm4f* | 1B | 5G | 0 | + |
| CP61* | Daghestan, Russia | VIR | WIR-10124 | *Pm4f* | 1-2B | 5G | 0 | + |
| CP73* | Daghestan, Russia | VIR | WIR-23924 | *Pm4f* | 1B | 5G | 0 | + |
| GR246* | Portugal | CGN | WAG6097 | *Pm4f* | 1B | 4-5G | 0 | + |
| IL50* | Turkey | NBRP | KU-10302 | *Pm4f* | 0 | 5G | 0 | + |
| IL131* | Turkey | NBRP | KU-10316 | *Pm4f* | 0 | 4G | 0 | + |
| GR341 | North Macedonia | CGN | CGN04172 | *Pm4f* | 1B | 2-3BG | 0-1B | + |
| IL186* | Armenia | NBRP | KU-1588 | *Pm4a* | 1B | 5G | 0 | + |
| CP20* | Azerbaijan | VIR | WIR-39311 | *Pm4a* | 0 | 4G | 0 | + |
| CP71* | Daghestan, Russia | VIR | WIR-23900 | *Pm4a* | 1B | 5G | 0 | + |
| IL92* | Iran | NBRP | KU-3282 | *Pm4a* | 0-1B | 5G | 0 | + |
| GR250* | Portugal | CGN | CGN06373 | *Pm4a* | 1B | 4G | 0-1B | + |
| CP27* | Tajikistan | VIR | WIR-24592 | *Pm4a* | 0 | 4G | 0 | + |
| CP30* | Tajikistan | VIR | WIR-24599 | *Pm4a* | 0-1B | 3-4G | 0 | + |
| IL132* | Turkey | NBRP | KU-10337 | *Pm4a* | 1B | 4G | 0-1B | + |
| GR192 | Germany | CRI | 01C010671 | *Pm4b* | 1B | 5G | 0-1B | + |
| GR130 | Austria | CGN | CGN05497 | *Pm4g* | 3G | 3G | 3G | - |
| GR142 | Austria | CGN | CGN14905 | *Pm4g* | 5G | 5G | 5G | - |
| GR176 | France | CGN | CGN04348 | *Pm4g* | 5G | 5G | 5G | - |
| GR180 | France | CGN | CGN05365 | *Pm4g* | 3G | 3G | 3G | - |
| GR210 | Germany | CGN | WAG5659 | *Pm4g* | 3-4G | 3-4G | 3-4G | - |
| CP81 | North Ossetia, Russia | VIR | WIR-14683 | *Pm4g* | 4G | 4G | 4G | - |
| IL16 | USA | NBRP | KU-306 | *Pm4g* | 3-4G | 4G | 4G | - |
| IL30 | Afghanistan | NBRP | KU-3074 | *PM4_h1* | 5G | 5G | 5G | - |
| IL233 | Iran | NBRP | KU-4315 | *PM4_h1* | 3G | 3-4G | 3-4G | - |
| IL234 | Iran | NBRP | KU-4335B | *PM4_h1* | 3G | 3-4G | 3-4G | - |

[a] Blue: Accessions reported to carry a single resistance gene at the *Rmg8* locus on 2BL through $F_2$ segregation analyses by Inoue et al. (2021) New Phytol. 229:488-500.

Yellow: Accessions subjected to $F_2$ segregation analyses in the present study. (See Extended Data Table 2.)

Asterisk: The 18 accessions reported to recognize *AVR-Rmg8* by Wang et al. (2018) Phytopathology 108:1299-1306.

[b] NBRP, National BioResource Project –Wheat, Japan (https://shigen.nig.ac.jp/wheat/komugi/strains/queryFormNbrp.jsp);
VIR, The N.I.Vavilov All-Russian Institute of Plant Genetic Resources (http://91.151.189.38/virdb/maindb);
JIC, John Innes Centre (https://www.seedstor.ac.uk/index.php);
CGN, Centre for Genetic Resources, the Netherlands (https://cgngenis.wur.nl/SearchDetails).
CRI, Crop Research Institute, Czech Republic

[c] 0 = no visible infection; 1 = pinhead spots; 2 = small lesions (<1.5 mm); 3 = scattered lesions of intermediate size (<3 mm); 4 = large typical lesions; and 5 = complete blighting of leaf blades. B and G represents brown and green lesions, respectively.

Resistant, moderately or weakly resistant, and susceptible interactions are shown in red, purple, and black, respectively.

Extended Data Table 2. Segregation of reactions to Br48 in $F_2$ populations derived from crosses between wheat lines.

| Cross[a] | Number of $F_2$ seedlings | | | $\chi2$ (15:1) | P |
|---|---|---|---|---|---|
| | Resistant | Susceptible | Total | | |
| S-615 (*Pm4f*) x IL186 (*Pm4a*) | 183 | 9 | 192 | 0.80 | 0.37 |
| S-615 (*Pm4f*) x CP20 (*Pm4a*) | 164 | 11 | 175 | 0.00 | 0.98 |

[a] *Rmg8* variants carried by the wheat lines are shown in parentheses.

Extended Data Table 3. Distribution of *Rmg8* variants in *Triticum* spp.

| Species | Region | Number of accessions | | | | | | | |
|---|---|---|---|---|---|---|---|---|---|
| | | *Pm4a* | *Pm4b* | *Pm4f* | *Pm4g* | *PM4_h1* | *PM4_h2* | Total of carriers | Total of accessions tested |
| *T. aestivum* | Total | 8 [1.5%] | 1 [0.2%] | 12 [2.3%] | 7 [1.3%] | 3 [0.6%] | 0 [0%] | 31 [5.9%] | 526 |
| | Europe, Middle East, and Africa | 7 | 1 | 12 | 6 | 3 | 0 | 29 | 336 |
| | Central Asia and Eastern Asia | 1 | 0 | 0 | 0 | 0 | 0 | 1 | 145 |
| | Americas | 0 | 0 | 0 | 1 | 0 | 0 | 1 | 45 |
| *T. durum* | Total | 4 [5.6%] | 0 [0%] | 2 [2.8%] | 0 [0%] | 0 [0%] | 0 [0%] | 6 [8.3%] | 72 |
| | Europe | 1 | 0 | 1 | 0 | 0 | 0 | 2 | 29 |
| | Middle East | 2 | 0 | 0 | 0 | 0 | 0 | 2 | 28 |
| | Ethiopia | 0 | 0 | 0 | 0 | 0 | 0 | 0 | 6 |
| | Central Asia and Eastern Asia | 1 | 0 | 0 | 0 | 0 | 0 | 1 | 5 |
| | Unknown | 0 | 0 | 1 | 0 | 0 | 0 | 1 | 4 |
| *T. dicoccum* | Total | 24 [31.6%] | 0 [0%] | 1 [1.3%] | 0 [0%] | 0 [0%] | 2 [2.6%] | 27 [35.5%] | 76 |
| | Europe Spain | 8 | 0 | 1 | 0 | 0 | 0 | 9 | 16 |
| | Others [a] | 2 | 0 | 0 | 0 | 0 | 0 | 2 | 25 |
| | Middle East | 0 | 0 | 0 | 0 | 0 | 0 | 0 | 12 |
| | Ethiopia | 8 | 0 | 0 | 0 | 0 | 2 | 10 | 12 |
| | India | 3 | 0 | 0 | 0 | 0 | 0 | 3 | 5 |
| | Unknown | 3 | 0 | 0 | 0 | 0 | 0 | 3 | 6 |
| *T. paleocolchicum* | Total | 0 [0%] | 0 [0%] | 0 [0%] | 0 [0%] | 0 [0%] | 0 [0%] | 0 [0%] | 4 |
| *T. dicoccoides* | Total [b] | 3 [6.5%] | 0 [0%] | 8 [17.4%] | 0 [0%] | 0 [0%] | 0 [0%] | 11 [23.9%] | 46 |

[a] Including accessions from Portugal, Italy, Belgium, Germany, Czech, Austria, Hungary, Bosnia and Herzegovina, Yugoslavia, Bulgaria, Romania, Russia, Armenia, Georgia, and Morocco.

[b] All accessions are from Fertile Crescent.

Extended Data Table 4. Reactions of representative accessions of *Ae. umbellulata*, *Ae. comosa*, and *Ae. speltoides* to Br48 and its *AVR-Rmg8* disruptant (Br48ΔA8)

| Species | Accession[a] | Infection type[b] with | | Recognition of |
|---|---|---|---|---|
| | | Br48 | Br48ΔA8 | *AVR-Rmg8* |
| *Ae. umbellulata* | KU-8-4 | 0-1B | 5G | + |
| | KU-8-5 | 0 | 5G | + |
| | KU-2737 | 1B | 5G | + |
| | KU-2738 | 0 | 5G | + |
| | KU-4002 | 1B | 5G | + |
| | KU-4005 | 1B | 5G | + |
| | KU-4006 | 1B | 5G | + |
| | KU-4009 | 0 | 5G | + |
| | KU-4013 | 0 | 5G | + |
| | KU-4014 | 0 | 5G | + |
| | KU-4018 | 0 | 5G | + |
| | KU-4026 | 0 | 5G | + |
| | KU-4035 | 0 | 5G | + |
| | KU-4036 | 0 | 5G | + |
| | KU-4040 | 0-1B | 3BG | + |
| | KU-4054 | 0 | 5G | + |
| | KU-4056 | 1B | 5G | + |
| | KU-4061 | 1B | 5G | + |
| | KU-4072 | 0 | 5G | + |
| | KU-4077 | 0 | 5G | + |
| | KU-4085 | 0-1B | 5G | + |
| | KU-4103 | 0 | 5G | + |
| | KU-4108 | 1B | 5G | + |
| | KU-5954 | 0 | 5G | + |
| | KU-12198 | 0 | 5G | + |
| | KU-12204 | 0 | 5G | + |
| | KU-12207a | 1B | 5G | + |
| | KU-4020 | 5G | 5G | - |
| | KU-4045 | 5G | 5G | - |
| | KU-12182 | 5G | 5G | - |
| | KU-12191 | 5G | 5G | - |
| *Ae. comosa* | KU-17-2 | 1B | 5G | + |
| *Ae. speltoides* | KU-2246 | 1B | 3G | + |
| | KU-2247 | 2B | 4G | + |
| | KU-2251 | 1B | 3G | + |
| | KU-7707 | 2B | 4G | + |
| | KU-7915 | 3BG | 5G | + |
| | KU-7921 | 1B | 5G | + |
| | KU-12962 | 2BG | 5G | + |
| | KU-2248 | 1B | 1B | - |
| | KU-2259 | 1B | 1B | - |
| | KU-12016 | 2B | 2B | - |

[a] All accessions that recognized *AVR-Rmg8* are shown with some examples that did not recognize *AVR-Rmg8*, i.e., four susceptible accessions of *Ae. umbellulata* used for crossing in Extended Data Table 5 and three resistant accessions of *Ae. speltoides* whose resistance was not impaired by the disruption of *AVR-Rmg8*.

[b] Refer to Extended Data Table 1. Red, resistant; purple, weakly resistant; black, susceptible.

Extended Data Table 5. Distribution of *Rmg8* variants in *Aegilops* spp.

| Species | Genome | Number of accessions | | |
|---|---|---|---|---|
| | | Resistant to Br48 | Recognizing *AVR-Rmg8* | Total |
| *Ae. umbellulata* | UU | 27 | 27 | 204 |
| *Ae. comosa* | MM | 1 | 1 | 4 |
| *Ae. speltoides* | SS | 18 | 7[a] | 139 |
| *Ae. tauschii* | DD | 13 | 0 | 204 |
| *Ae. caudata* | CC | 2 | 0 | 247 |
| *Ae. uniaristata* | NN | 0 | 0 | 11 |
| *Ae. longissima* | $S^lS^l$ | 2 | 0 | 33 |
| *Ae. sharonensis* | $S^{sh}S^{sh}$ | 4 | 0 | 15 |
| *Ae. searsii* | $S^sS^s$ | 0 | 0 | 19 |
| *Ae. bicornis* | $S^bS^b$ | 0 | 0 | 33 |
| Total | - | 67 | 35 | 909 |

[a] Among the 18 accessions resistant to Br48, eight accesssions have not been tested with Br48ΔA8 because sufficient seeds were not available. The seven accessions recognizing *AVR-Rmg8* were detected in the other 10 accessions with sufficient seeds.

Extended Data Table 6. Segregation of reactions to Br48 in $F_2$ populations derived from crosses between *Aegilops umbellulata* accessions.

| Cross[a] | Number of $F_2$ seedlings | | | $\chi^2$ (3:1) | P |
|---|---|---|---|---|---|
| | Resistant | Susceptible | Total | | |
| KU-4013 (R) x KU-12182 (S) | 29 | 9 | 38 | 0.04 | 0.85 |
| KU-4013 (R) x KU-4045 (S) | 116 | 34 | 150 | 0.44 | 0.51 |
| KU-4014 (R) x KU-12182 (S) | 32 | 8 | 40 | 0.53 | 0.47 |
| KU-4014 (R) x KU-4045 (S) | 81 | 31 | 112 | 0.43 | 0.51 |
| KU-4036 (R) x KU-12182 (S) | 26 | 12 | 38 | 0.88 | 0.35 |
| KU-4036 (R) x KU-4045 (S) | 30 | 8 | 38 | 0.32 | 0.57 |
| KU-2737 (R) x KU-4020 (S) | 28 | 8 | 36 | 0.15 | 0.70 |
| KU-4054 (R) x KU-4020 (S) | 34 | 6 | 40 | 2.13 | 0.14 |
| KU-4085 (R) x KU-12191 (S) | 29 | 8 | 37 | 0.23 | 0.64 |
| KU-4013 (R) x KU-4014 (R) | 70 | 0 | 70 | – | – |
| KU-4036 (R) x KU-4014 (R) | 73 | 0 | 73 | – | – |
| KU-2737 (R) x KU-4054 (R) | 39 | 0 | 39 | – | – |
| KU-4054 (R) x KU-4085 (R) | 33 | 0 | 33 | – | – |

[a] Reactions of parental accessions to Br48 are shown in parentheses. R, resistant; S, susceptible.

Extended Data Table 7. Reactions of *Triticum aestivum* lines and *Aegilops umbellulata* accessions to wheat powdery mildew isolates collected in Japan.

| | | | Infection type[a] on | | | | | | | | | | |
|---|---|---|---|---|---|---|---|---|---|---|---|---|---|
| | | | *T. aestivum* [b] | | | | | | | *Ae. umbellulata* [b] | | | |
| | | | N4 | ChI | ChI-Rmg8 | Cc | Cc-Pm4 | IL16 | GR142 | KU-4026 | KU-4013 | KU-4020 | KU-4043 |
| Isolates | Year | Location | (-) | (-) | (*Pm4f*) | (-) | (*Pm4a*) | (*Pm4g*) | (*Pm4g*) | (*AeuRmg8_h1*) | (nd) | (nd) | (nd) |
| Th1 | 2005 | Matsusaka City, Mie | 4 | 4 | 4 | 4 | 4 | 4 | 4 | 4 | 4 | 4 | 4 |
| Th2 | 2021 | Kyoto City, Kyoto | 4 | 4 | 4 | 4 | 0 | 4 | 4 | 4 | 3 | 4 | 4 |
| Th3 | 2021 | Chikugo City, Fukuoka | 4 | 4 | 4 | 4 | 4 | 3 | 3 | 4 | 3 | 4 | 4 |
| Th4 | 2021 | Chikugo City, Fukuoka | 4 | 4 | 4 | 4 | 4 | 4 | 4 | 4 | 3 | 4 | 4 |
| Th5 | 2021 | Ichikawa-cho, Hyogo | 4 | 4 | 4 | 4 | 4 | 4 | 4 | 4 | 3 | 4 | 4 |
| Th6 | 2021 | Kasai City, Hyogo | 4 | 4 | 4 | 4 | 4 | 4 | 4 | 3 | 3 | 3 | 3 |
| Th7 | 2021 | Tsukuba City, Ibaraki | 4 | 4 | 4 | 4 | 4 | 4 | 4 | 4 | 3 | 4 | 4 |
| Th8 | 2021 | Tsukuba City, Ibaraki | 4 | 4 | 4 | 4 | 4 | 4 | 4 | 4 | 3 | 4 | 4 |
| Th9 | 2021 | Tsukuba City, Ibaraki | 4 | 4 | 4 | 4 | 4 | 4 | 4 | 4 | 4 | 4 | 4 |
| Th10 | 2021 | Tsukuba City, Ibaraki | 4 | 4 | 4 | 4 | 4 | 4 | 3 | 4 | 4 | 4 | 4 |
| Th12 | 2021 | Kobe City, Hyogo | 4 | 4 | 4 | 4 | 4 | 4 | 4 | 4 | 3 | 4 | 4 |
| Th13 | 2021 | Muko City, Kyoto | 4 | 4 | 4 | 4 | 4 | 4 | 4 | 3 | 3 | 3 | 4 |
| Th14 | 2021 | Muko City, Kyoto | 4 | 4 | 4 | 4 | 4 | 4 | 4 | 4 | 4 | 4 | 4 |
| Th15 | 2021 | Obihiro City, Hokkaido | 4 | 4 | 4 | 4 | 0 | 4 | 4 | 4 | 4 | 4 | 4 |

[a] 0, no mycelial growth or sporulation; 1, scant sporulation; 2, reduced sporulation; 3, slightly reduced sporulation; 4, heavy sporulation. Incompatible interactions are shown in red.

[b] Lines/accessions resistant and susceptible to MoT isolate Br48 are painted pink and green, respectively. *Rmg8* variants carried by them are shown in parentheses. nd, not determined.

Extended Data Table 8. Primers used in this study

| Objectives | Primer name | Sequence( 5' -> 3') | Description |
|---|---|---|---|
| Mapping of *Rmg8* | KM9-F | TGATGGTCTGACGTCCGTGC | CAPS marker (*Spe* I) |
| | KM9-R | CAATCACTACCAGTAACGTTACACGGTG | |
| | KM12-F | TAGAGCTCTCAAGCACTTCCCTTGAG | CAPS marker (*Hinf* I) |
| | KM12-R | GAAGTGGAAAGGTTAGACACTGAGGAAG | |
| | KM13-F | GTGCCACCGAGGCGACTTGTTC | CAPS marker (*Msp* I) |
| | KM13-R | AGTCTCCCCCAAAGTTCCAGCGG | |
| | KM25-F | AGGTAGATGCTGCTATGTGACTTGTG | Presence/absence marker (presence in Sch) |
| | KM25-R | AGACGGAGATTGTGATGAGGAG | |
| | KM27-F | AGGCACGGAAGGCAATTTCAC | Presence/absence marker (presence in S-615) |
| | KM27-R | TGCTCGCTGATAGCATCAACAAG | |
| | KM30-F | AACCGTGAGATTTCCACTGC | Presence/absence marker (presence in S-615) |
| | KM30-R | TATCTCCGTAAAGAGCTTCCAAGA | |
| | KM65-F | ACGATGCATCCTTGATACATCAAC | CAPS marker (*Hae* III) |
| | KM65-R | TCCGTGTCACAGTTCAGAAATAGATAG | |
| | KM109-F | TGGATTTGTTCTTCATCGCTTC | CAPS marker (*Hae* III) |
| | KM109-R | TGTTGGAACGTAGGGTTCACC | |
| | KM138-F | ATAACGCTGAGTCAAAGTCTCCAC | Presence/absence marker (presence in S-615) |
| | KM138-R | TAGATACGGTTGAAATGCTTCTTCTC | |
| | KM140-F | AGCATCATCCAAAAAGTCCC | Presence/absence marker (presence in S-615) |
| | KM140-R | TGTATTGAGAAATCGTTACCTGG | |
| | KM155-F | AGATGGGGATTGACAGCTTG | Presence/absence marker (presence in Sch) |
| | KM155-R | AGAACATGTGCCACTACAAGGC | |
| Mapping of *Rmg8* & Detection of *Rmg8* variants | KM171-F | TGACGCTAGAGACCAACAGATG | Presence/absence marker (presence in S-615) |
| | KM171-R | ACCATTGGAAGGATGAGCTG | |
| Mapping of *Rmg8* and *Rmg7* & Detection of Rmg8 variants | KM200-F | TTCGACGGCATCTGCAAGTGGAAGAACCC | Presence/absence marker (presence in S-615) |
| | KM200-R | ACTGCGCGCGCTCCCCCTGCAT | |
| Mapping of *Rmg7* | KM201-F | AAAAATAGAGGTGCGTGGGTAG | CAPS marker (*Mbo* I) |
| | KM201-R | AAAAGAGGGGGATTGAAGGAG | |
| | KM202-F | AAGGGTCAGGCGTTAATGG | CAPS marker (*Xho* I) |
| | KM202-R | ATCCAGCATCCTGCACATTT | |
| Transgene detection* | HPT-F | GTGTCACGTTGCAAGACCTG | for detection of transgene |
| | HPT-R | GATGTTGGCGACCTCGTATT | |
| Protoplast assay & transformation of Fielder | InF-Rmg8-V1-F | TGTGTGTGCAGATCGGGTTCTACTGACTGCAAATG | for In-Fusion cloning of *Rmg8-V1* and *Rmg7-V1* |
| | InF-Rmg8-V1-R | GGAAATTCGAGCTCGAACCACATTTCACAAGAGAG | |
| | InF-Rmg8-V2-F | TGTGTGTGCAGATCGGGTTCTACTGACTGCAAATG | for In-Fusion cloning of *Rmg8-V2* and *Rmg7-V2* |
| | InF-Rmg8-V2-R | GGAAATTCGAGCTCGCGACGTCATCAATACTTCAT | |
| | InF-Rmg8-genome-F | TGTGTGTGCAGATCGGGTTCTACTGACTGCAAATG | for In-Fusion cloning of *Rmg8* -genome |
| | InF-Rmg8-genome-R | GGAAATTCGAGCTCGCGACGTCATCAATACTTCAT | |
| Protoplast assay | InF-PWT3dSP-F | TGTGTGTGCAGATCGATGAGTGACTTTTGGAAAGTA | for In-Fusion cloning of *PWT3* without its signal peptide |
| | InF-PWT3dSP-R | GGAAATTCGAGCTCGTTACGGCGATGCAAACAGC | |
| | InF-AVR-Rmg8dSP-F | TGTGTGTGCAGATCGATGCTGCCTGCGCCGCAG | for In-Fusion cloning of *AVR-Rmg8* without its signal peptide |
| | InF-AVR-Rmg8dSP-R | GGAAATTCGAGCTCGCTACTGCCTTCTAGTACCG | |
| *Rmg8* variant analysis | Rmg8-E5-F | GGTTCTACTGACTGCAAATGGA | for amplification of Exon 1-5 of *Rmg8* variants in *Triticum* spp. and *Ae. speltoides* from genomic DNA |
| | Rmg8-E5-R | TGTAGCAACCCAATTAAAGGAAG | |
| | Rmg8-V1V2-F | GAAACGTGCTACCAGACAGAATC | for amplification of Exon 6 (V1) and Exon 7 (V2) of *Rmg8* variants in *Triticum* spp. and *Ae. speltoides* from genomic DNA |
| | Rmg8-V1V2-R | CAACATGAAGAAATTCATCGTCA | |
| | Rmg8U-V1-1R | ATTTCACAAGAGAGCTAGCGG | for amplification of *AeuRmg8* V1 variant from cDNA of *Ae. umbellulata* accessions in combination with Rmg8-E5-F |
| | Rmg8U-V2-1R | AAATTCATCATCACAGGAGCAC | for amplification of *AeuRmg8* V2 variant from cDNA of *Ae. umbellulata* accessions (KU-4026, KU-4103, KU-5934, KU-12198) in combination with Rmg8-E5-F |
| | Rmg8U-V2-2R | AAGTAATTGCAACATGAAGAAATTC | for amplification of *AeuRmg8* V2 variant from cDNA of *Ae. umbellulata* accessions (KU-8-5, KU-4035, KU-4043, KU-5954) in combination with Rmg8-E5-F |
| | Rmg8M-V1-1R | GTCAGGTCAGCAGGTGG | for amplification of *AecRmg8* V1 variant from cDNA of *Ae. comosa* (KU-17-2) with Rmg8-E5-F |
| | Rmg8M-V2-1R | AAATTCATCGTCACAGGAGC | for amplification of *AecRmg8* V2 variant from cDNA of *Ae. comosa* (KU-17-2) with Rmg8-E5-F |
| | M13_fwd | GTAAAACGACGGCCAGT | for sanger sequencing |
| | T3_promoter | ATTAACCCTCACTAAAGGGAA | for sanger sequencing |
| | Rmg8-seq-F1 | GAAACCCGCCAATATACTGCTC | for sanger sequencing |
| | Rmg8-seq-F2 | ATGCTATGAGACAGCTTTAGAGTGC | for sanger sequencing |
| | Rmg8-seq-F3 | GAAACGTGCTACCAGACAGAATC | for sanger sequencing |
| | Rmg8-seq-F4 | AAGTGGTTCAGCCTGTTGGTG | for sanger sequencing |
| | Rmg8-seq-F5 | ATGACGCTAGAGACCAACAGATG | for sanger sequencing |
| | Rmg8-seq-F6 | AGTATGGCAAACCGCAGCTC | for sanger sequencing |
| | Rmg8-seq-F7 | ATCCTTCCAATGGTGTTACTGTG | for sanger sequencing |
| | Rmg8-seq-R1 | TCCATAAGAGCATTGGACATCC | for sanger sequencing |
| | Rmg8-seq-R2 | TTTGTCCGGTCAGTATTTCTAGG | for sanger sequencing |
| | Rmg8-seq-R3 | CACGACCTTGAGGTACATGAGTT | for sanger sequencing |
| | Rmg8-seq-R4 | TCCGTCTCAAGGCTCAGATG | for sanger sequencing |
| | Rmg8-seq-R5 | GGACATGAAGCGGTAGAAGTTG | for sanger sequencing |
| | Rmg8-seq-R6 | TGTTGTCCATCTTGTGGAGC | for sanger sequencing |
| | Rmg8-seq-R7 | CAACATGAAGAAATTCATCGTCA | for sanger sequencing |
| | Rmg8U_V1_seq_F1 | ACCTTCATGATAATCATATTATT | for sanger sequencing |
| | Rmg8U_V1_seq_R1 | TTGGCGGGTTTTAGATCCAAGTG | for sanger sequencing |
| | Rmg8U_V2_seq_F1 | GCCGTGAAAAGGATGTCCAATGC | for sanger sequencing |
| | Rmg8U_V2_seq_F2 | AGCGCTGACTTGTATAGTCTTGG | for sanger sequencing |
| | Rmg8U_V2_seq_R1 | AGCGGCGGGTCGGCCTGCTCCAG | for sanger sequencing |
| | Rmg8U_V2_seq_R2 | CGAGCTCGGCGTGTGACAGCGTC | for sanger sequencing |

* Refer to Abe, F. et al. Genome-edited triple-recessive mutation alters seed dormancy in wheat. *Cell Rep* . **28**, 1362-1369 (2019).
